# MeCP2 is Necessary in Cerebellar Purkinje Cells for Precise Network Dynamics During Associative Motor Learning

**DOI:** 10.64898/2026.05.25.727684

**Authors:** Jenny Shen, Peter Shen, Julia Lopes Gonçalez, Mikayla Jackson, Likhitha Polepalli, Kruthika Dheeravath, Timothy Li, Abdulla Hamki, Mariam Hamki, Christian Hicks, Chang Li, Lucas Pozzo-Miller, Wei Li

## Abstract

Loss-of-function variants of *MECP2* cause Rett syndrome; however, their impact on cerebellar computations for learning remains poorly understood. Here, we show that *Mecp2* deletion specifically in Purkinje cells does not broadly disrupt cerebellar-dependent behaviors but selectively compromises those that require precise timing, coordination, and associative updating in the cerebellar cortex. *Mecp2* loss altered the emergence of learning-related Purkinje cell activity *in vivo* and disrupted their intrinsic and synaptic properties that support adaptive cerebellar output. These findings identify MeCP2 as a critical player in Purkinje cell function to regulate cerebellar learning signals and suggest that Rett syndrome-related motor dysfunction reflects impaired adaptive computation rather than a generalized loss of motor capacity.

## Introduction

Rett syndrome (RTT) is a severe X-linked neurodevelopmental disorder caused in most cases by loss-of-function variants of *methyl-CpG-binding protein 2* (*MECP2*), a transcription regulator that is essential for neuronal maturation and circuit stability^1, 2^. After an initial period of apparently typical development, affected individuals undergo regression marked by loss of acquired speech and purposeful hand use, emergence of repetitive behaviors, intellectual disability, and progressive motor impairment^3–5^. Motor dysfunction is one of the most disabling features of RTT and includes altered gait, ataxia, apraxia, hypotonia, and deterioration of learned movements^6^. Mouse models with global or CNS loss of *Mecp2* similarly develop RTT-like phenotypes, including impaired motor coordination and motor learning, supporting the view that brain dysfunction is a major driver of disease pathophysiology^7–10^.

In addition to the forebrain, the most extensively studied brain region in RTT, converging neuropathological, imaging, and experimental studies point to an important contribution from the cerebellum^11^. Neuropathological studies in RTT have reported reduced cerebellar size together with atrophy of the molecular and granule cell layers^12,13^. Consistent with these human findings, *Mecp2*-deficient mouse models also show reduced brain volume and altered cerebellar morphology^14, 15^. These observations are especially notable because the cerebellum is essential for timing, coordination, prediction and motor learning^16, 17^, and is now understood to participate in broader cerebello-thalamo-cortical and cerebello-basal ganglia networks that shape cognitive, affective and social behaviors^18–21^. Thus, cerebellar dysfunction is well positioned to contribute to both the motor and non-motor manifestations of RTT^22^.

Two recent studies have begun to address how *Mecp2* loss in the cerebellum affects behavior, but they point to distinct and still unresolved mechanisms. Achilly and colleagues showed that deleting *Mecp2* from the cerebellum causes a delay in motor learning that can be overcome with additional training, whereas deletion from individual cerebellar neuronal subtypes, including Purkinje cells (PCs), did not reproduce the same phenotype^23^; notably, these mice showed preserved social behaviors but irregular firing in PCs. In contrast, Xu et al. reported that PC-specific *Mecp2* deficiency is sufficient to impair motor learning and produce autistic-like behaviors and further linked these phenotypes to altered intrinsic excitability of PCs^24^. Together, these studies strongly implicate the cerebellum, and especially PCs, in RTT pathophysiology, but they also leave unresolved how *Mecp2* loss in PCs reshapes learning-related cerebellar activity across behavioral, circuit and cellular levels.

These unresolved issues are important because PCs are the sole output neurons of the cerebellar cortex and occupy a strategic position for transforming cellular alterations into circuit-level deficits in timing, error signaling, and adaptive motor control^16, 18^. It remains unclear whether PC-specific *Mecp2* deficiency selectively disrupts only a narrow set of cerebellar learning paradigms or instead perturbs a broader hierarchy of motor behaviors, from gait coordination to skilled forelimb movement to associative learning. It is also unknown how PC-specific *Mecp2* loss alters learning-related activity *in vivo* at the levels of single-cell firing and population dynamics, or how these changes relate to dendritic remodeling, altered intrinsic excitability, excitatory synaptic organization, and altered plasticity within the cerebellar cortex.

Here, we combined cell-type-specific genetic deletion, quantitative behavioral analyses, *in vivo* electrophysiology and fiber photometry of fluorescent sensors, morphological reconstructions, and *ex vivo* intracellular recordings to define how *Mecp2* deletion in PCs perturbs cerebellar function across multiple levels of organization. We show that loss of *Mecp2* in PCs leaves gross locomotor and social behaviors largely intact but impairs cerebellum-dependent motor learning and disrupts learning-related PC activity. At the cellular level, *Mecp2*-deficient PCs exhibit structural, intrinsic, and synaptic abnormalities that identify them as a critical locus through which *Mecp2* loss impairs cerebellar learning. More broadly, these findings support the idea that RTT motor phenotypes do not arise solely from cortical or basal ganglia dysfunction but also reflect atypical computation within cerebellar output neurons.

## Results

### PC-specific *Mecp2* deletion preserves general behaviors

We generated PC-specific *Mecp2* conditional knockout (cKO) mice by crossing *Pcp2*-Cre mice^25^ with *Mecp2* floxed mice^8^, and we used both female homozygous (*Pcp2*-Cre; *Mecp2*^fl/fl^) and male hemizygous (*Pcp2*-Cre; *Mecp2*^fl/y^) mice, along with their wildtype (WT) littermates as controls. To confirm PC-specific deletion of *Mecp2*, we examined MeCP2 immunoreactivity in cerebellar sections co-labeled for calbindin (CB) for PCs, S100β for astrocytes, and parvalbumin (PV) for GABAergic interneurons. In *Mecp2* cKO mice, MeCP2 immunoreactivity was selectively absent from CB-positive PCs but remained in S100β-positive astrocytes and PV-positive interneurons (Fig. 1a); MeCP2 was detected in all three cell types in control WT mice (Extended Data Fig. 1a).

**Fig. 1.**
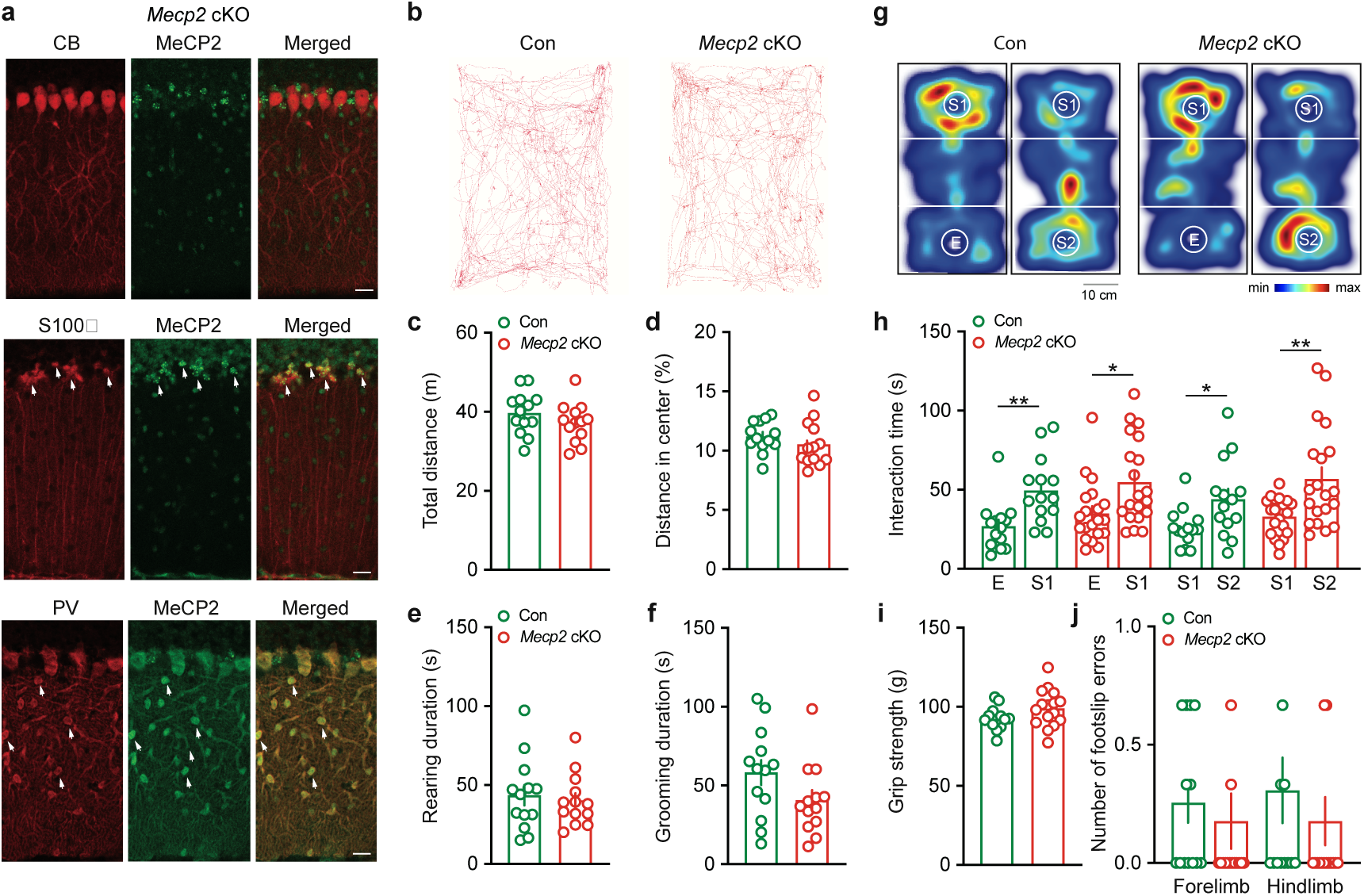
| Behavioral and cellular characterization of *Mecp2* cKO mice. **a,** Immunofluorescence of cerebellar sections showing the PC marker calbindin (CB), the astrocyte marker S100β, and the interneuron marker parvalbumin (PV) with MeCP2 labeling, confirming loss of MeCP2 expression in PCs of *Mecp2* cKO mice. Arrows indicate colocalization. Scale bar, 30 μm. **b,** Representative locomotor trajectories in the open field for control and *Mecp2* cKO mice. **c,d,** Total distance traveled in the entire arena (**c**) and distance traveled in the center zone relative to total distance (**d**). Unpaired two-sided Student’s *t*-test (n = 13, Con; n = 13, *Mecp2* cKO; *P* = 0.2218, total distance; *P* = 0.2344, center distance). **e,f,** Rearing (**e**) and grooming (**f**) durations during open field testing. Unpaired two-sided Student’s *t*-test (n = 13, Con; n = 13, *Mecp2* cKO; *P* = 0.6786, rearing duration; *P* = 0.0940, grooming duration). **g,** Heat maps showing spatial occupancy of control and *Mecp2* cKO mice encountering the empty inverted cup (E) or cups containing novel mice (S1 or S2) during the three-chamber social test. Minimal and maximum values indicate the relative accumulating activity in each area of the arena. **h,** Time spent interacting with E, S1, or S2 (n = 14, Con; n = 20, *Mecp2* cKO). Unpaired two-sided Student’s *t*-test (*P* = 0.0030, Con E vs. S1; *P* = 0.0139, *Mecp2* cKO E vs. S1; *P* = 0.0220, Con S1 vs. S2; *P* = 0.0032, *Mecp2* cKO S1 vs. S2). **i,** Forelimb grip strength in control (n = 13) and *Mecp2* cKO (n = 16) mice. Unpaired two-sided Student’s *t*-test (*P* = 0.0968, Con vs. *Mecp2* cKO). **j,** Foot-slip errors of forelimbs and hindlimbs on horizontal ladder rungs in control (n = 13) and *Mecp2* cKO (n = 15) mice. Mann-Whitney two-sided test (*P* = 0.2031, forelimb; *P* = 0.3706, hindlimb; Con vs. *Mecp2* cKO). All data are presented as mean ± SEM. Individual points represent individual animals.

Using these animals, we next assessed general behavioral function. In the open field, *Mecp2* cKO mice were indistinguishable from controls in total distance traveled, velocity, and immobility (Fig. 1b,c and Extended Data Fig. 1b,c). They also showed normal center-zone distance, center-zone time, and center-zone entries, suggesting that PC-specific *Mecp2* deletion does not increase anxiety-like behaviors (Fig. 1d and Extended Data Fig. 1d,e). Rearing and grooming duration and frequency were also unchanged, consistent with preserved exploration and spontaneous self-directed behaviors (Fig. 1e,f and Extended Data Fig. 1f,g).

We next assessed social behaviors using the classical three-chamber assay (Fig. 1g). Both groups spent more time interacting with the novel social stimulus (stranger 1, S1) than with the empty cup (E) in the sociability phase, and with a second novel mouse (S2) than with the first novel mouse (S1) in the social memory phase, indicating intact social preference and short-term social memory in *Mecp2* cKO mice (Fig. 1h). To control for possible effects of locomotor activity on interaction time, we calculated a preference index by normalizing the difference in interaction time to the total interaction time; this measure did not differ between groups (Extended Data Fig. 1h). Forelimb grip strength and ladder-rung performance were also comparable between genotypes, with no increase in forelimb or hindlimb foot-slip errors in *Mecp2* cKO mice (Fig. 1i,j). Together, these data indicate that PC-specific *Mecp2* deletion does not produce broad deficits in locomotor activity, anxiety-like behavior, social preference and memory, or gross motor function.

### PC-specific *Mecp2* deletion causes subtle gait coordination deficits

Because standard behavioral assays did not reveal major abnormalities in general motor function, we next asked whether PC-specific *Mecp2* deletion produces subtler deficits in locomotor coordination. Consistent with the open-field, grip-strength, and ladder-rung results, automated *CatWalk* gait analysis showed that several basic gait parameters were preserved in *Mecp2* cKO mice (Fig. 2a). Stride length and step cycle were comparable between genotypes across all four paws (Fig. 2b,c), and stand time, swing time, swing speed and print area were also largely unchanged (Extended Data Fig. 2a–d). The overall distribution of step-sequence patterns was also not grossly altered (Extended Data Fig. 2e,f), arguing against a major disruption of basic stepping organization.

**Fig. 2.**
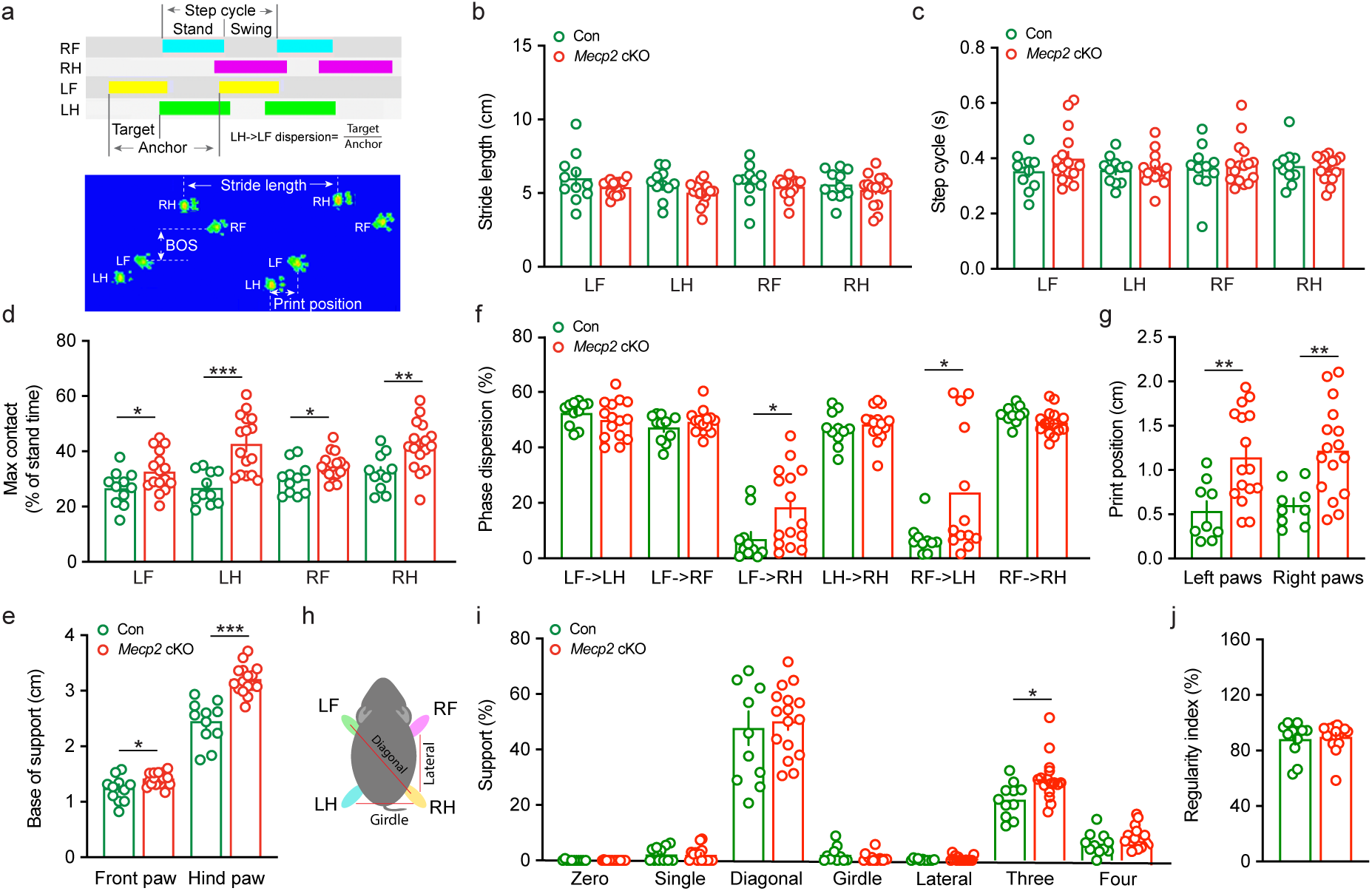
| Gait and interlimb coordination analysis of *Mecp2* cKO mice. **a**, Schematic representation of gait parameters measured using the CatWalk system. The step cycle is divided into stand and swing phases for each limb: left front (LF), left hind (LH), right front (RF), and right hind (RH). Stride length is defined as the distance between consecutive placements of the same paw, and base of support (BOS) represents the lateral distance between left and right paws. Phase dispersion reflects the temporal coordination between limb pairs and is calculated as the timing of initial contact of a target paw relative to the step cycle of an anchor paw, expressed as a percentage. Print position reflects the distance between forepaw and the subsequent placement of the ipsilateral hind paw. Representative footprint heatmap illustrates paw placement and stride pattern. **b,** Stride length for each limb (LF, LH, RF, RH) in control and *Mecp2* cKO mice. Unpaired two-sided Student’s *t*-test (n = 11, Con; n = 16, *Mecp2* cKO; *P* = 0.1910, LF; *P* = 0.0802, LH; *P* = 0.4064, RF; *P* = 0.3710, RH). **c,** Step cycle duration for each limb (LF, LH, RF, RH). Unpaired two-sided Student’s *t*-test (n = 11, Con; n = 16, *Mecp2* cKO; *P* = 0.1916, LF; *P* = 0.9523, LH; *P* = 0.6013, RF; *P* = 0.6979, RH). **d,** Maximum contact time, expressed as a percentage of stand time, for each limb (LF, LH, RF, RH). Unpaired two-sided Student’s *t*-test (n = 11, Con; n = 16, *Mecp2* cKO; *P* = 0.0347, LF; *P* = 0.0001, LH; *P* = 0.0416, RF; *P* = 0.0054, RH). **e,** BOS for front and hind paws. Unpaired two-sided Student’s *t*-test (n = 11, Con; n = 16, *Mecp2* cKO; *P* = 0.0160, front paws; *P* < 0.001, hind paws). **f,** Phase dispersion between limb pairs (LF→LH, LF→RF, LF→RH, LH→RH, RF→LH, RF→RH), reflecting interlimb coordination. Unpaired two-sided Student’s *t*-test or Mann–Whitney two-sided test (n = 11, Con; n = 15, *Mecp2* cKO; *P* = 0.2977, LF→LH; *P* = 0.2694, LF→RF; *P* = 0.0246, LF→RH; *P* = 0.5212, LH→RH; *P* = 0.0255, RF→LH; *P* = 0.0969, RF→RH). **g,** Print position for left and right paws. Unpaired two-sided Student’s *t*-test (n = 9, Con; n = 16, *Mecp2* cKO; *P* = 0.0029, left paws; *P* = 0.0045, right paws). **h,** Schematic of support patterns used to classify gait coordination, including diagonal, lateral, and girdle configurations. **i,** Distribution of support patterns (zero, single, diagonal, girdle, lateral, three-paw, and four-paw support), expressed as percentage of total steps. Unpaired two-sided Student’s *t*-test or Mann-Whitney two-sided test (n = 11, Con; n = 16, *Mecp2* cKO; *P* = 0.7493, zero; *P* = 0.8771, single; *P* = 0.6903, diagonal; *P* = 0.3091, girdle; *P* = 0.9682, lateral; *P* = 0.0149, three; *P* = 0.114, four). **j,** Regularity index, representing the percentage of normal step sequence patterns. Unpaired two-sided Student’s *t*-test (n = 11, Con; n = 16, *Mecp2* cKO; *P* = 0.6906). All data are presented as mean ± SEM. Individual points represent individual animals.

By contrast, more refined analyses of interlimb coordination uncovered a selective locomotor phenotype. Maximal contact timing was altered across multiple paws (Fig. 2d), indicating a shift in the time point at which each paw reached peak surface contact during stance. Because maximal contact reflects how the body weight is transferred onto the paws, this finding suggests impaired stance-phase control rather than reduced strength. Consistently, base of support (BOS) was increased, particularly in the hind paws (Fig. 2e), indicating that *Mecp2* cKO mice adopted a wider stance, likely as a compensatory strategy to enhance stability.

Additional alterations pointed to impaired temporal and spatial coordination among limbs. Phase dispersion was altered for limb pairs (Fig. 2f), suggesting that the relative timing between paw placements was less precisely controlled in *Mecp2* cKO mice. Because phase dispersion captures the temporal relationship between target and anchor limbs during the step cycle, these changes are most consistent with a deficit in fine interlimb coupling rather than global slowing of movement. Print position was also shifted for both left and right paws (Fig. 2g), indicating abnormal placement of hind paws relative to preceding forepaw positions and therefore reduced spatial precision of sequential stepping. In parallel, paw-support configuration was redistributed (Fig. 2h,i), with greater use of three-paw support states. Because support patterns reflect how animals transition between stable and unstable phases of gait, these findings further support the presence of subtle instability during locomotion. Notably, the regularity index remained largely unchanged (Fig. 2j), indicating that overall step sequence consistency was preserved. Together, these results indicate that PC-specific *Mecp2* deletion leaves gross motor behavior largely intact but disrupts higher-order features of gait that depend on precise cerebellar coordination.

### PC-specific *Mecp2* deletion differentially affects motor learning across behavioral paradigms

To determine whether PC-specific *Mecp2* deletion affects motor learning, we compared performance across skilled reaching, rotarod, and eyeblink conditioning assays, which engage partially overlapping but distinct neural circuits. In the single-pellet reaching task (Fig. 3a), which depends strongly on corticospinal and motor cortical control of forelimb kinematics^26, 27^, *Mecp2* cKO mice showed largely preserved task acquisition. Mice were trained for 20 min per day over 12 days to retrieve pellets with the preferred forelimb, and both groups increased total reaching attempts across training (Fig. 3c). Success rate also improved over days in both groups, with no significant difference between the two genotypes in learning trajectory (Fig. 3d). To quantify forelimb kinematics, we reconstructed three-dimensional reaching trajectories from dual-camera recordings and analyzed individual digits, the forepaw and the pellet using *DeepLabCut*-based tracking (Fig. 3b). Total reach and grasp distance and duration were not significantly altered across training (Fig. 3e,f), and separate analysis of reach distance, grasp distance, reach time and grasp time similarly revealed no major genotype-dependent differences (Extended Data Fig. 3a-d). However, *Mecp2* cKO mice showed greater reach deviation across training, while grasp deviation was unchanged (Fig. 3g,h), suggesting that PC-specific *Mecp2* deletion subtly affects forelimb trajectory precision without broadly disrupting skilled reaching acquisition or the overall structure of the movement.

**Fig. 3.**
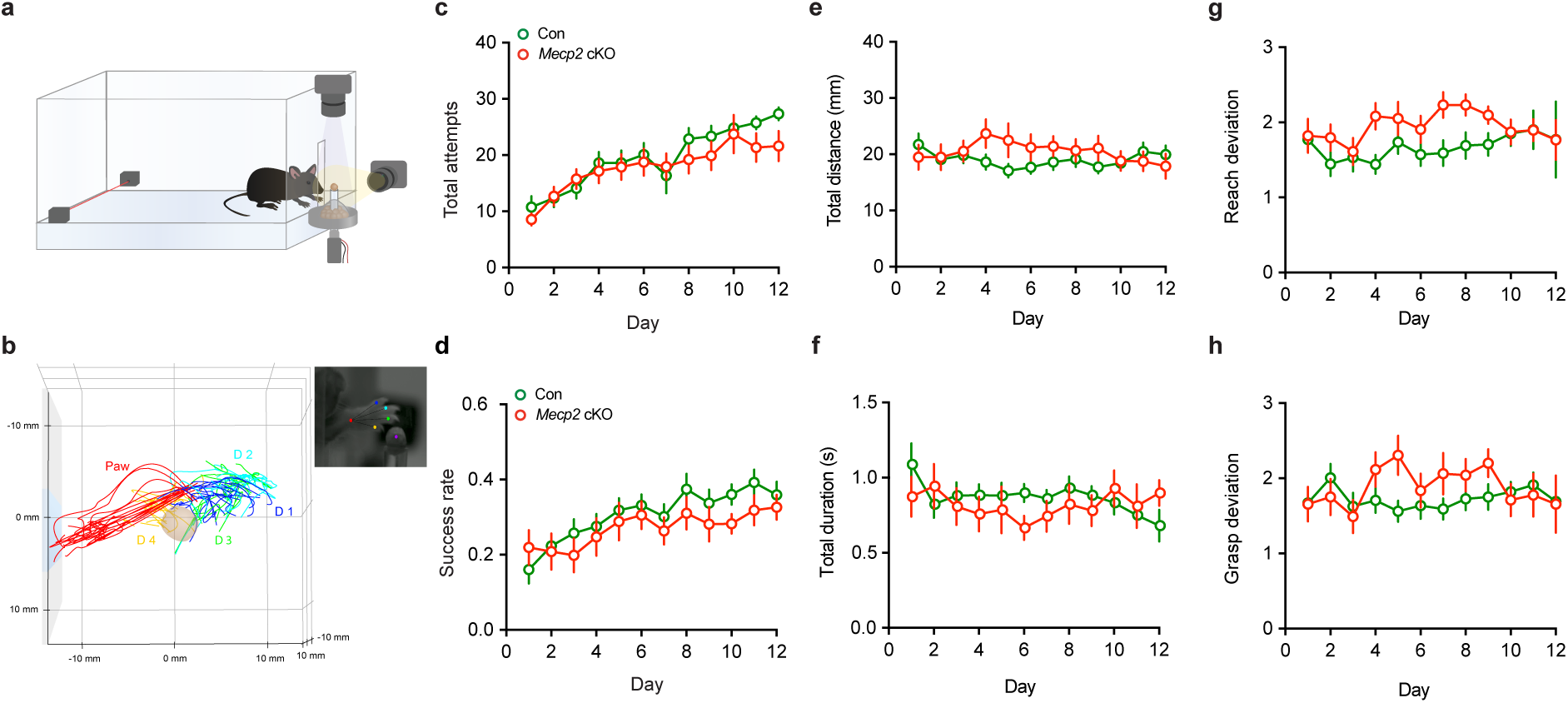
| Skilled reaching performance in *Mecp2* cKO mice. **a**, Schematic of the automated single-pellet reaching task with dual-camera recording for three-dimensional kinematic analysis. **b,** Representative three-dimensional forelimb trajectories reconstructed from recordings and analyzed using DeepLabCut-based tracking of individual digits, the forepaw, and the pellet. **c,** Total reaching attempts across training days. Two-way repeated-measures ANOVA showed no session × genotype interaction *F*(11,384) = 1.178, *P* = 0.3002), a significant main effect of day *F*(11,384) = 20.86, *P* < 0.0001, and no main effect of genotype *F*(1,35) = 0.3355, *P* = 0.5662 (n = 13, Con; n = 24, *Mecp2* cKO). **d,** Success rate across training days. Two-way repeated-measures ANOVA showed no session × genotype interaction *F*(11,242) = 0.7458, *P* = 0.6849, a significant main effect of day *F*(11,242) = 6.689, *P* < 0.0001, and no main effect of genotype *F*(1,22) = 1.038, *P* = 0.3194 (n = 14, Con; n = 10, *Mecp2* cKO). **e,** Total reach and grasp distance across training days. Two-way repeated-measures ANOVA showed no session × genotype interaction *F*(11,383) = 1.014, *P* = 0.4332, no main effect of day *F*(11,383) = 0.4815, *P* = 0.9146, and no main effect of genotype *F*(1,51) = 0.9410, *P* = 0.3366 (n = 27, Con; n = 26, *Mecp2* cKO). **f,** Total reach and grasp duration across training days. Two-way repeated-measures ANOVA showed no session × genotype interaction *F*(13,397) = 1.520, *P* = 0.1070, no main effect of day *F*(13,397) = 0.4832, *P* = 0.9338, and no main effect of genotype *F*(1,51) = 0.05047, *P* = 0.9944 (n = 25, Con; n = 28, *Mecp2* cKO). **g,** Reach deviation across training days. Two-way repeated-measures ANOVA showed no session × genotype interaction *F*(13,415) = 1.405, *P* = 0.1534, a significant main effect of day *F*(13,415) = 2.175, *P* = 0.0099), and a significant main effect of genotype *F*(1,51) = 8.341, *P* = 0.0057 (n = 25, Con; n = 28, *Mecp2* cKO). **h,** Grasp deviation across training days. Two-way repeated-measures ANOVA showed no session × genotype interaction *F*(13,398) = 1.326, *P* = 0.1945, no main effect of day *F*(13,398) = 1.460, *P* = 0.1296), and no main effect of genotype *F*(1,51) = 0.1444, *P* = 0.7055 (n = 25, Con; n = 28, *Mecp2* cKO). All data are presented as mean ± SEM.

We next tested motor learning on the rotarod, a task that depends heavily on cerebellar coordination and adaptation to an accelerating rotating rod^28^. In contrast to the largely preserved pellet-reaching performance, *Mecp2* cKO mice showed a shorter latency to fall compared with controls, with significant deficits emerging during later training days (Extended Data Fig. 4a).

We then asked whether associative motor learning was affected by assessing eyeblink conditioning (Fig. 4a). In delay eyeblink conditioning, the conditioned stimulus (CS) and unconditioned stimulus (US) overlap each other, and associative learning depends predominantly on cerebellar circuitry (Fig. 4b,c). *Mecp2* cKO mice showed reduced conditioned response (CR) acquisition during the early phase of training, but gradually improved and approached control levels later in training (Fig. 4d,e); eyelid closure amplitude followed a similar delayed acquisition profile. Thus, although associative motor learning was slowed, repeated training was sufficient to partially compensate for the deficit in the delay paradigm. Conversely, deficits were more pronounced in trace eyeblink conditioning, in which a stimulus-free interval separates the CS from the US, and successful learning requires not only cerebellar processing but also coordinated engagement of other brain regions, including the hippocampus and prefrontal cortex^29^. In this paradigm, *Mecp2* cKO mice showed persistently reduced CR percentage and reduced eyelid closure amplitude during the 12-day acquisition period, with deficits remaining evident during the 2-day reacquisition phase (Fig. 4f,g). Thus, whereas delay conditioning was slowed down but ultimately partially recovered to control levels, trace conditioning remained substantially impaired, indicating that PC-specific *Mecp2* deletion has a stronger impact when the task places greater demands on distributed learning-related networks and on the temporal bridging of sensory events. These data suggest that loss of *Mecp2* in PCs preferentially compromises forms of motor learning that rely heavily on cerebellar timing, coordination, and associative plasticity, particularly when these processes must be integrated with broader cognitive circuitry.

**Fig. 4.**
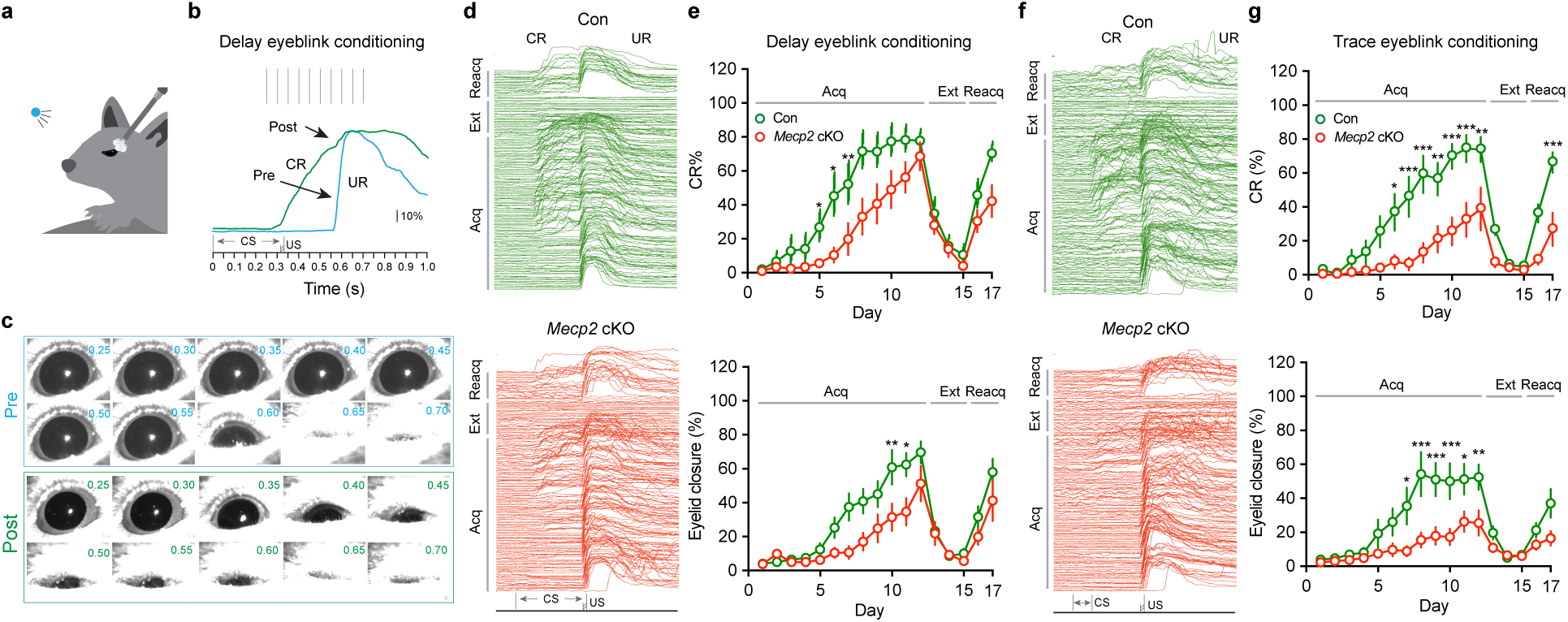
| Impaired delay and trace eyeblink conditioning in *Mecp2* cKO mice. **a**, Schematic of the head-fixed eyeblink conditioning setup. **b,** Representative eyelid trace during delay eyeblink conditioning before and after training, illustrating CRs and URs. In delay eyeblink conditioning, the CS was a 350-ms blue LED light and the US was a 30-ms air puff, with the CS and US co-terminating. **c,** Representative eye images during delay eyeblink conditioning before and after training, corresponding to the blue traces in **b**. **d,e,** Representative eyelid traces (**d**) and quantification of CR percentage (**e**) across acquisition (Acq), extinction (Ext), and reacquisition (Reacq) during delay eyeblink conditioning. Mice underwent 12 acquisition sessions, 3 extinction sessions, and 2 reacquisition sessions. For CR percentage, two-way repeated-measures ANOVA with Bonferroni’s multiple-comparisons test showed a significant session ξ genotype interaction *F*(16,288) = 2.474, *P* = 0.0015, a significant main effect of session *F*(16,288) = 37.03, *P* < 0.0001, and a significant main effect of genotype *F*(1,18) = 6.639, *P* = 0.0190 (n = 10, Con; n = 10, *Mecp2* cKO). Post hoc testing identified significant differences on acquisition days 6, 7, and 8 (*P* < 0.05). For eyelid closure percentage, two-way repeated-measures ANOVA with Bonferroni’s multiple-comparisons test showed a significant session ξ genotype interaction *F*(16,288) = 2.526, *P* = 0.0012, a significant main effect of session *F*(16,288) = 26.26, *P* < 0.0001, and a significant main effect of genotype *F*(1,18) = 7.097, *P* = 0.0158). Post hoc testing identified significant differences on acquisition days 10 and 11 (*P* < 0.05). **f,g,** Representative eyelid traces (**f**) and quantification of CR percentage (**g**) across acquisition, extinction, and reacquisition during trace eyeblink conditioning. In trace eyeblink conditioning, the CS was a 50-ms light stimulus followed by a 270-ms trace interval and then a 30-ms air puff US. For CR percentage, two-way repeated-measures ANOVA with Bonferroni’s multiple-comparisons test showed a significant session ξ genotype interaction *F*(16,240) = 5.004, *P* < 0.0001, a significant main effect of session *F*(16,240) = 27.20, *P* < 0.0001, and a significant main effect of genotype *F*(1,15) = 16.72, *P* = 0.0010 (n = 8, Con; n = 9, *Mecp2* cKO). Post hoc testing identified significant differences on acquisition days 6-12 and reacquisition day 2 (*P* < 0.05). For eyelid closure percentage, two-way repeated-measures ANOVA with Bonferroni’s multiple-comparisons test showed a significant session ξ genotype interaction *F*(16,240) = 4.131, *P* < 0.001, a significant main effect of session *F*(16,240) = 17.88, *P* < 0.001), and a significant main effect of genotype *F*(1,15) = 8.160, *P* = 0.012. Post hoc testing identified significant differences on acquisition days 7-10 and day 12 (*P* < 0.05). All data are presented as mean ± SEM. Individual points represent individual animals.

### Altered learning-related PC firing during trace eyeblink conditioning

To examine how PC activity is modulated during associative learning, we performed acute *in vivo* single-unit recordings from PCs in head-fixed mice during trace eyeblink conditioning (Fig. 5a). Recordings were obtained during early acquisition (EA; days 1-2) and late acquisition (LA; days 9-12), representing pre-learning and post-learning stages, while eyelid movement, treadmill movement, and electrophysiological signals were synchronized. Representative traces and raster plots show PC simple spikes (SmSpks) and complex spikes (CxSpks) aligned to the onset of CS and US in control mice (Fig. 5b,c). Averaged firing profiles further indicate that learning was accompanied by a prominent reorganization of event-related PC firing, most notably for CxSpks (Fig. 5d,e). We first asked whether conditioning altered SmSpk output. Although clear SmSpk responses were detected around CS and US presentation, and individual PCs often showed transient suppression after the US (Fig. 5c), quantification of CS-aligned and US-aligned normalized SmSpk responses did not reveal significant differences between EA and LA in either genotype (Extended Data Fig. 5a,b). Thus, under these conditions, PC SmSpk output did not show a significant learning-dependent change at the population level. In contrast, CxSpk responses were modulated in a learning-dependent manner and differed between genotypes. In control mice, CS-aligned normalized CxSpk responses increased from EA to LA, pointing to the emergence of a learning-related CxSpk signal as training progressed (Fig. 5f). Meanwhile, US-aligned normalized CxSpk responses were reduced at LA relative to EA, indicating a redistribution of CxSpk signaling across the course of learning (Fig. 5g). In *Mecp2* cKO mice, however, this learning-associated reorganization was disrupted. *Mecp2* cKO mice failed to show the same increase in CS-aligned CxSpk activity or the same shift in US-aligned responses seen in controls. These findings indicate that PC-specific *Mecp2* deletion impairs the normal learning-related reorganization of CxSpk signaling during trace eyeblink conditioning.

**Fig. 5.**
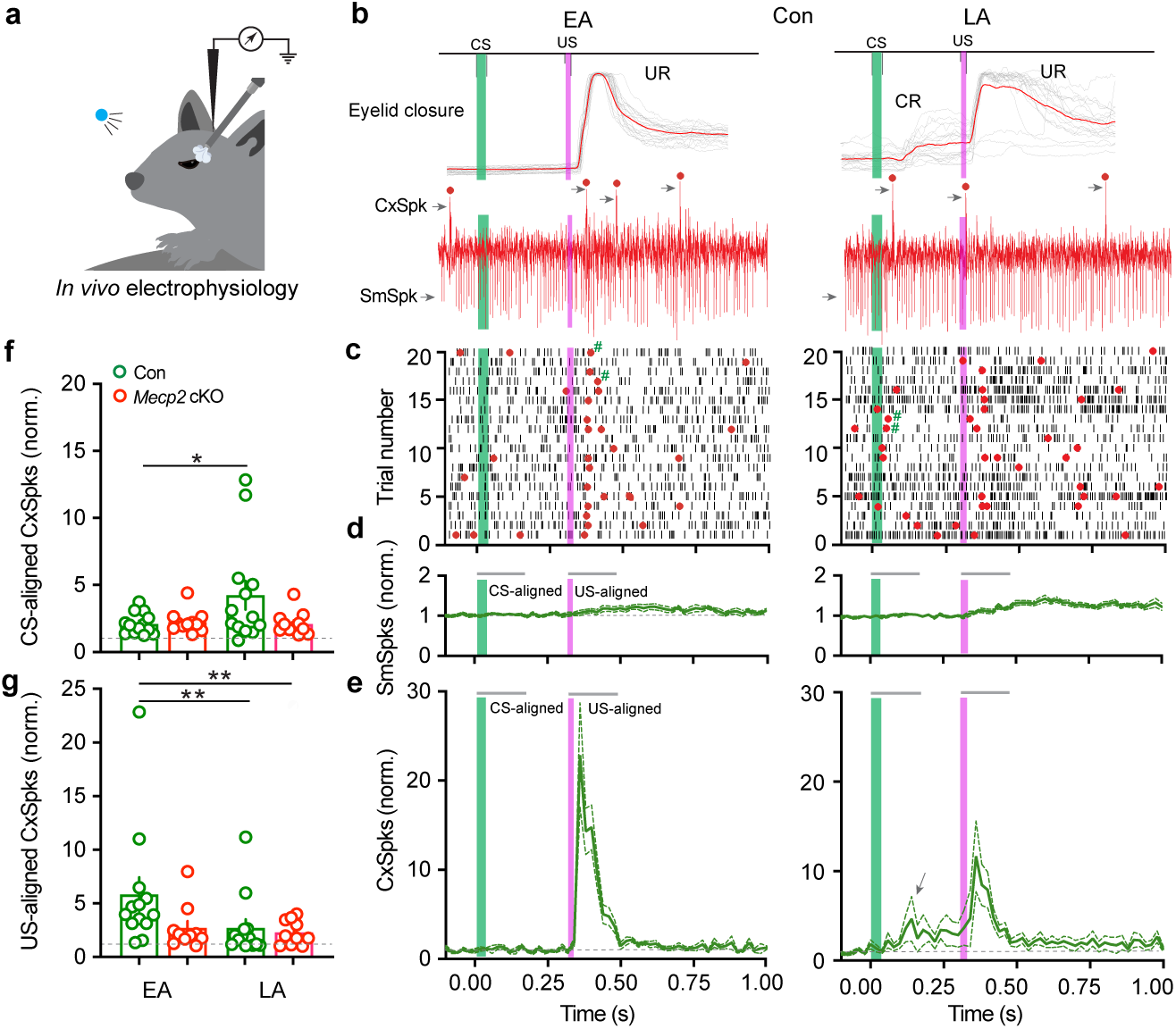
| Altered PC SmSpk and CxSpk responses during eyeblink conditioning in *Mecp2* cKO mice. **a**, Schematic of the head-fixed *in vivo* electrophysiology setup during trace eyeblink conditioning. Acute extracellular recordings were obtained during early acquisition (EA; days 1-2) and late acquisition (LA; days 9-12) of eyeblink conditioning. Eyelid movement, treadmill movement, and electrophysiological signals were synchronized using TTL pulses. **b,** Representative eyelid closure traces and simultaneously recorded PC activity during EA and LA in control mice. SmSpks (arrowheads) and CxSpks (red dots and arrowheads) are aligned to conditioned stimulus (CS) and unconditioned stimulus (US) onset. **c,** Representative raster plots of PC firing aligned to CS and US onset during EA and LA in control and *Mecp2* cKO mice. Red dots indicate CxSpks, short black lines indicate SmSpks, and # indicates the inhibition of SmSpks following CxSpks. **d,e,** Averaged SmSpk firing profiles (**d**) and CxSpk firing profiles (**e**) aligned to CS and US onset during EA and LA. Firing rates were calculated in 20-ms bins and normalized to baseline activity. Arrows indicate the increase in CxSpks following eyeblink conditioning. **f,** Quantification of CS-aligned normalized CxSpk responses in control and *Mecp2* cKO mice during EA and LA. Wilcoxon matched-pairs signed-rank test (n = 13, Con; n = 13, *Mecp2* cKO; *P* = 0.0398, Con EA vs. Con LA). **g,** Quantification of US-aligned normalized CxSpk responses in control and *Mecp2* cKO mice during EA and LA. Mann-Whitney two-sided test (n = 13, Con; n = 13, *Mecp2* cKO; *P* = 0.0086, Con EA vs. Con LA; *P* = 0.0099, Con EA vs. *Mecp2* cKO LA). All data are presented as mean ± SEM. Individual points represent individual animals.

To determine whether these firing changes simply reflected behavioral performance, we next examined the relationship between spike responses and CR percentage. Across EA and LA, neither CS– nor US-aligned SmSpk responses correlated significantly with CR percentage in either genotype (Extended Data Fig. 5c,d). Likewise, CS– and US-aligned CxSpk responses did not show significant correlations with CR percentage (Extended Data Fig. 5e,f). Therefore, the altered firing patterns in *Mecp2* cKO mice are unlikely to be explained solely by trial-by-trial differences in learned eyelid behavior.

Because locomotor state can influence cerebellar activity, we also assessed running speed during trace eyeblink conditioning. Running-speed profiles differed between CS-aligned and US-aligned epochs and across learning phases, indicating that animals changed locomotor state during task performance. In both control and *Mecp2* cKO mice, running speed was consistently lower during the US-aligned epoch than during the CS-aligned epoch in both the EA and LA phases (Extended Data Fig. 5g-i). However, normalized running speed did not correlate significantly with CS– or US-aligned SmSpk or CxSpk responses in either genotype during either EA or LA (Extended Data Fig. 5j-m). These results argue that the altered PC firing observed in *Mecp2* cKO mice is not secondary to changes in locomotion. Together, these single-cell results show that PC-specific *Mecp2* deletion selectively disrupts learning-related modulation of CxSpk activity during trace eyeblink conditioning, while largely sparing SmSpk responses. This impairment in adaptive CxSpk signaling suggests that loss of *Mecp2* in PCs interferes with cerebellar instructive or error-related signals that normally emerge during associative motor learning^30, 31^.

### Altered learning-related population Ca^2+^ signals in PCs during trace eyeblink conditioning

We further determined how population-level PC activity evolves during associative learning by performing *in vivo* fiber photometry of intracellular Ca^2+^ signals as surrogates for spiking activity in GCaMP6f-expressing PCs during trace eyeblink conditioning (Fig. 6a). Bilateral fiber-optic cannulas were implanted over ipsilateral and contralateral lobule VI relative to the air puff (US), and Ca^2+^ signals were recorded throughout acquisition, extinction, and reacquisition, while eyelid behavior and treadmill movement were synchronized. In mice with cannula implants, the impairment in eyeblink conditioning in *Mecp2* cKO mice remained evident (Extended Data Fig. 6a,b). For each trial, Ca^2+^ signals were extracted from –1.0 to 2.0 s relative to stimulus onset, baseline-subtracted, z-scored, and aligned to CS or US onset (Fig. 6b, *top*); trials associated with locomotor bout initiation were excluded from averaged analyses (Fig. 6b, *middle* and *bottom*). Representative CS– and US-aligned traces in control mice showed prominent learning-related modulation of PC population Ca^2+^ signals between EA and LA (Fig. 6b), and heat maps across acquisition, extinction, and reacquisition further indicated that these activity patterns evolved gradually, with responses becoming progressively more prominent during the CS period and less dominant during the US period as CRs emerged, diminished, and reappeared (Fig. 6c).

**Fig. 6.**
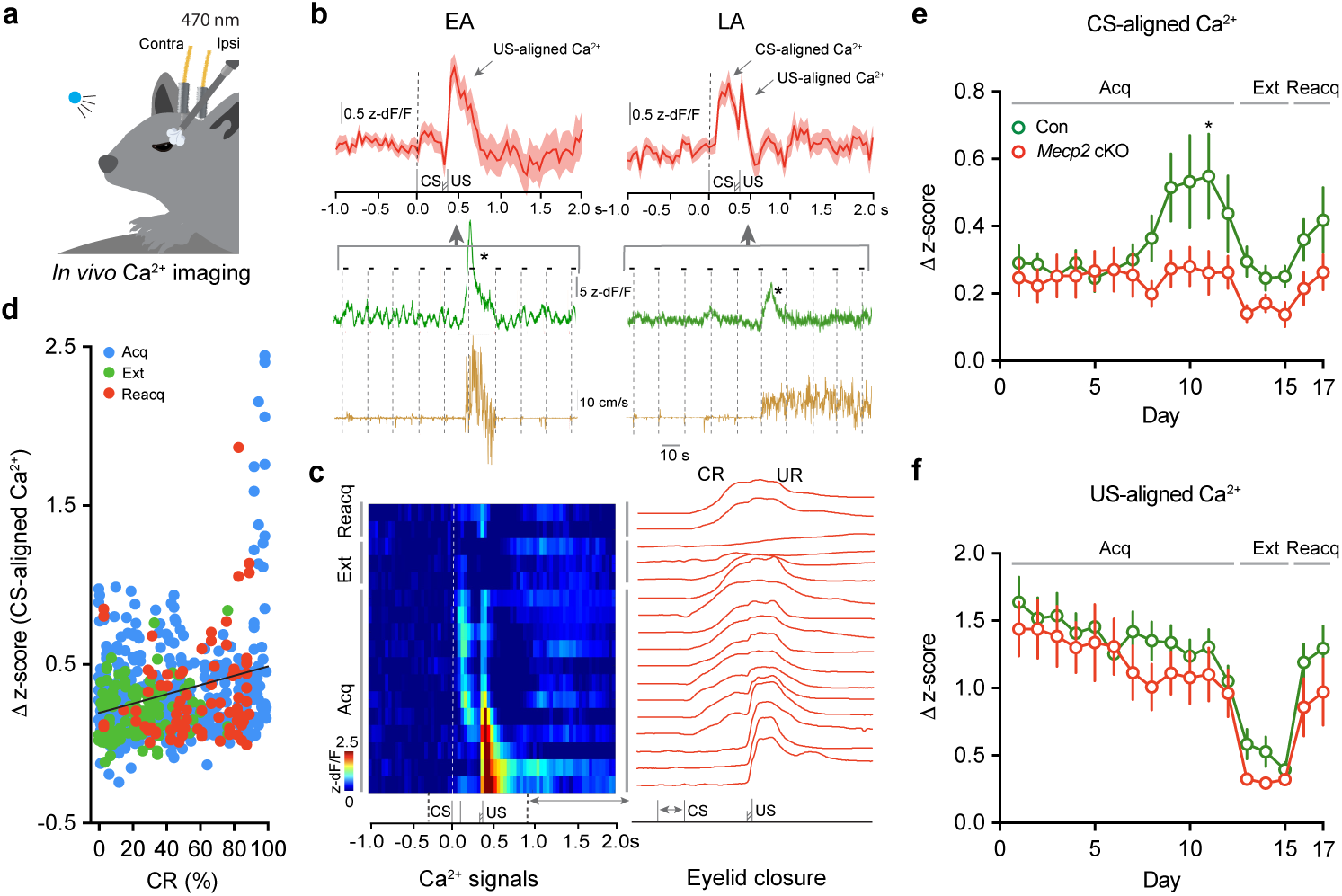
| Learning-related PC Ca^2+^ signals during trace eyeblink conditioning in *Mecp2* cKO mice. **a**, Schematic of the head-fixed *in vivo* Ca^2+^ photometry setup during trace eyeblink conditioning. GCaMP6f signals were recorded bilaterally from PCs through implanted fiber-optic cannulas positioned over the ipsilateral and contralateral cerebellar lobule VI. **b,** Representative CS– and US-aligned Ca^2+^ traces and running-speed traces during EA and LA in control mice. Ca^2+^ signals were extracted within an analysis window from –1.0 to 2.0 s relative to stimulus onset, averaged across all air-puff trials (short lines), and expressed as z-scored dF/F. Trials associated with the onset of locomotor rotation bouts within 2 s (asterisk in example trace) were excluded from averaged analyses. **c,** Heat maps of session-by-session CS– and US-aligned Ca^2+^ activity across acquisition (Acq), extinction (Ext), and reacquisition (Reacq), with representative eyelid closure traces from the same behavioral phases. **d,** Relationship between CS-aligned Ca^2+^ responses and CR percentage across acquisition, extinction, and reacquisition. Each point represents the session-averaged Ca^2+^ response plotted against behavioral performance (CR percentage) from all control and *Mecp2* cKO animals. Simple linear regression showed a significant correlation between CR percentage and z-scored dF/F (n = 680 session-averaged Ca^2+^ signals; R^2^ = 0.07149, *F*(1,678) = 52.21, *P* < 0.001). For event-related analyses, peak Ca^2+^ responses were quantified within defined post-stimulus windows relative to baseline activity. **e,f,** Quantification of CS-aligned Ca^2+^ responses (**e**) and US-aligned Ca^2+^ responses (**f**) across acquisition, extinction, and reacquisition in control and *Mecp2* cKO mice. For CS-aligned Ca^2+^ signals, two-way repeated-measures ANOVA with Bonferroni’s multiple-comparisons test showed a significant day ξ genotype interaction *F*(16,608) = 1.815, *P* = 0.0261, a significant main effect of day *F*(16,608) = 3.656, *P* < 0.001, and a trend toward significant main effect of genotype *F*(1,38) = 3.253, *P* = 0.0792 (n = 21, Con; n = 19, *Mecp2* cKO). Post hoc testing identified a significant difference on day 11 (*P* = 0.0338). For US-aligned Ca^2+^ signals, two-way repeated-measures ANOVA with Bonferroni’s multiple-comparisons test showed no significant day ξ genotype interaction *F*(16,608) = 0.6189, *P* = 0.8700, a significant main effect of day *F*(16,608) = 27.74, *P* < 0.001, and no significant main effect of genotype *F*(1,38) = 0.9749, *P* = 0.3297 (n = 21, Con; n = 19, *Mecp2* cKO). All data are presented as mean ± SEM. Individual points represent individual animals.

Across all sessions in both genotypes, CS-aligned Ca^2+^ responses were positively correlated with CR percentage, indicating that stronger learning-related PC population Ca^2+^ signals were associated with better behavioral performance (Fig. 6d). Across training, CS-aligned population Ca^2+^ signals in control mice gradually increased during acquisition, peaked late in training, declined during extinction, and rose again during reacquisition. In *Mecp2* cKO mice, this learning-related increase was blunted, resulting in weaker CS-evoked PC population activity across training (Fig. 6e). By contrast, US-aligned Ca^2+^ responses gradually decreased during acquisition, diminished further during extinction and recovered during reacquisition in control mice, and a similar pattern was observed in *Mecp2* cKO mice (Fig. 6f), indicating that the population-level abnormality in *Mecp2* cKO mice was more profound in the CS-related than in the US-related component of the signal.

To further define the relationship between PC Ca^2+^ activity and behavior, we examined phase-specific correlations with CR percentage during acquisition, extinction, and reacquisition. During acquisition, CS-aligned Ca^2+^ responses correlated significantly with CR percentage in control mice but not in *Mecp2* cKO mice, indicating that the typical coupling between learning-related PC population activity and behavioral performance was weakened by *Mecp2* deletion in PCs (Extended Data Fig. 6c). US-aligned Ca^2+^ responses during acquisition also correlated with CR percentage in both genotypes, although this relationship was weaker than that seen for CS-aligned signals in controls (Extended Data Fig. 6d). In contrast, neither CS– nor US-aligned responses correlated significantly with CR percentage during extinction or reacquisition in either genotype, indicating that the tightest relationship between PC population Ca^2+^ dynamics and behavior occurred during initial learning (Extended Data Fig. 6e-h).

Because Ca^2+^ recordings were obtained bilaterally, we separated the signals into ipsilateral and contralateral hemispheres and next asked whether these population responses were lateralized. Learning-related CS– and US-aligned Ca^2+^ responses were strongest ipsilateral to the US air puff but were also detectable contralaterally, suggesting that eyeblink conditioning recruits distributed cerebellar activity rather than a strictly unilateral local circuit. The genotype-dependent difference was most apparent ipsilaterally, where control mice developed a stronger training-related increase in population Ca^2+^ responses, whereas this increase was reduced in *Mecp2* cKO mice (Extended Data Fig. 6i,j). However, both ipsilateral and contralateral US-aligned Ca^2+^ responses followed similar trajectories across training in control and *Mecp2* cKO mice (Extended Data Fig. 6k,l). These findings suggest that the genotype-dependent disruption in learning-related PC population activity was most evident in ipsilateral CS-aligned signals rather than in US-aligned responses.

Together, these data show that PC-specific *Mecp2* deletion disrupts learning-dependent population Ca^2+^ dynamics during trace eyeblink conditioning, most prominently in CS-aligned signals, suggesting that loss of *Mecp2* in PCs impairs the learning-dependent recruitment of PC ensembles during associative motor learning.

### Structural and synaptic remodeling of PCs in *Mecp2* cKO mice

To determine whether the altered cerebellar PC activity and associative learning deficits in *Mecp2* cKO mice were accompanied by structural changes in PCs, we filled individual PCs with biocytin during intracellular whole-cell recordings in *ex vivo* cerebellar slices, and examined dendritic architecture, dendritic spine features, and markers of excitatory synapses (Fig. 7a,b). Sholl analyses of dendritic morphology revealed reduced dendritic complexity in *Mecp2* cKO PCs, with fewer dendritic intersections at distal radii from the soma, indicating a simplified dendritic arbor (Fig. 7c); consistently, overall dendritic coverage area was also smaller in *Mecp2* cKO mice (Fig. 7d). We next asked whether dendritic spines were altered (Fig. 7e). The numerical density of dendritic spines was unchanged, but spine volume was larger in *Mecp2* cKO mice (Fig. 7f,g), indicating that PC-specific *Mecp2* deletion remodels existing postsynaptic structures without causing a detectable loss of dendritic spine numbers.

**Fig. 7.**
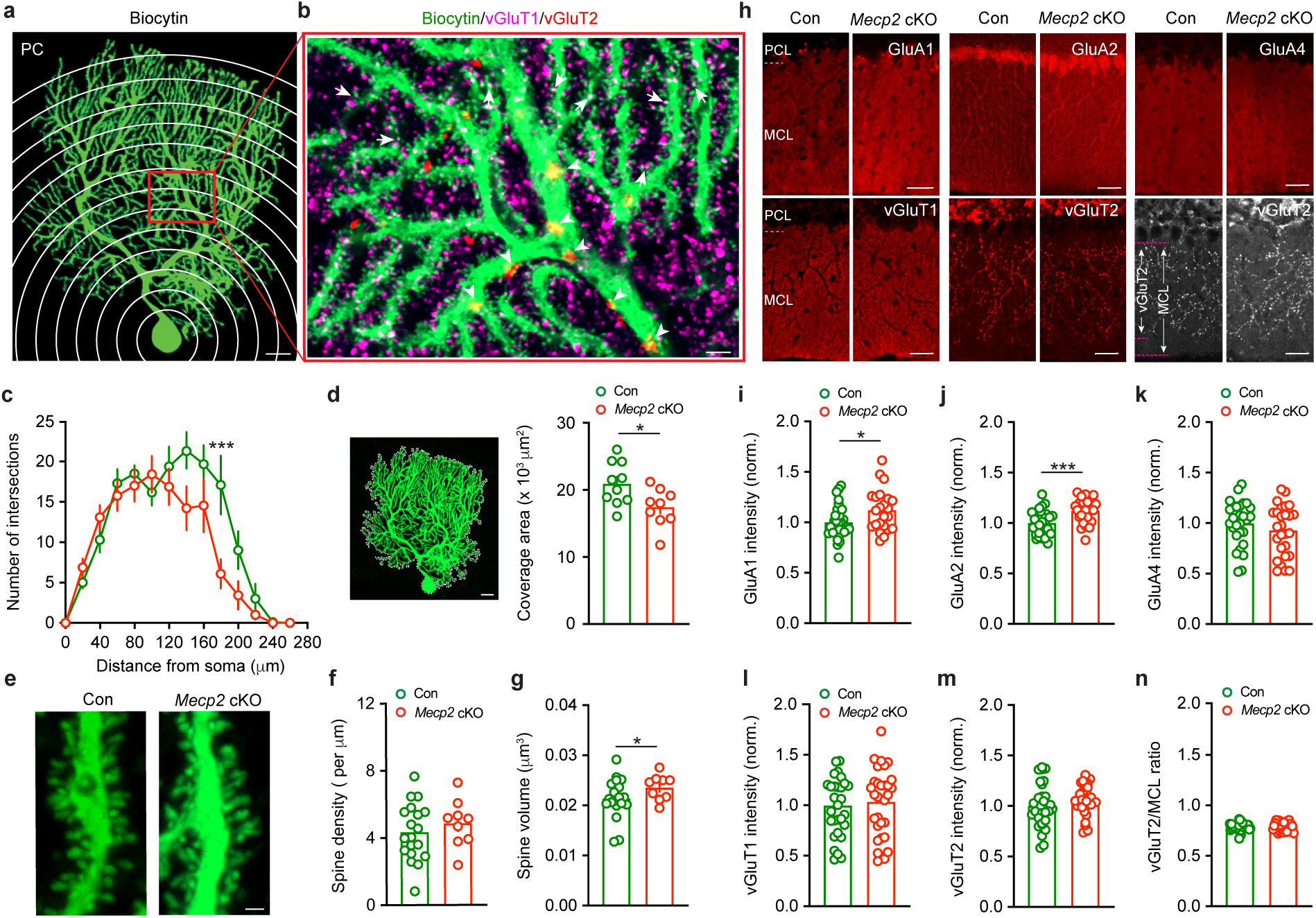
| Structural and synaptic alterations in PCs in *Mecp2* cKO mice. **a,b**, PCs were filled with biocytin during intracellular whole-cell recordings in *ex vivo* cerebellar slices, followed by vGluT1 and vGluT2 immunostaining. Three-dimensional Sholl analysis was used to assess the number of dendritic intersections. Arrows indicate colocalization of vGluT1 puncta (magenta) with biocytin-filled PC dendritic spines (green), whereas arrows indicate vGluT2 puncta (red) on PC dendrites. Scale bars, 20 μm (**a)** and 10 μm (**b)**. **c,** The number of intersections with 3D Sholl spheres centered at the PC soma is plotted as a function of distance from the soma. Two-way repeated-measures ANOVA with Bonferroni’s multiple-comparisons test showed a significant distance ξ genotype interaction, *F*(13, 221) = 2.43, *P* = 0.0044, and significant main effects of distance *F*(13, 221) = 35.86, *P* < 0.001 and genotype *F*(1, 17) = 4.60, *P* = 0.0467 (n = 10, Con; n = 9, *Mecp2* cKO). Post hoc testing identified a significant difference at 180 μm (*P* < 0.001). **d,** PC coverage was measured from the outline of the PC. Unpaired two-sided Student’s *t*-test (n = 10, Con; n = 9, *Mecp2* cKO; *P* = 0.0212). Scale bar, 10 μm. **e,** Representative images of biocytin-filled PC dendritic spines. Scale bar, 2 μm. **f,g,** Quantification of average spine density (**f**) and spine volume (**g**). Unpaired two-sided Student’s *t*-test (n = 19, Con; n = 9, *Mecp2* cKO; *P* = 0.1722, spine density; *P* = 0.0453, spine volume). **h,** Immunostaining for GluA1, GluA2, GluA4, vGluT1, and vGluT2 in the cerebellar Purkinje cell layer (PCL) and molecular cell layer (MCL), with an example showing measurements of vGluT2 extension and MCL length. Scale bar, 30 μm. **i-m,** Quantification of GluA1 (**i**, n = 27, Con; n = 24, *Mecp2* cKO), GluA2 (**j**, n = 27, Con; n = 24, *Mecp2 cKO*), GluA4 (**k**, n = 28, Con; n = 27, *Mecp2* cKO), vGluT1 (**l**, n = 29, Con; n = 31, *Mecp2 cKO*), and vGluT2 (**m**, n = 32, Con; n = 31, *Mecp2 cKO*) immunoreactivity in *Mecp2* cKO mice normalized to control mice. Unpaired two-sided Student’s *t*-test (*P* = 0.0231, GluA1; *P* < 0.001, GluA2; *P* = 0.2549, GluA4; *P* = 0.6746, vGluT1; *P* = 0.2100, vGluT2). **n,** Quantification of the vGluT2/MCL length ratio. Two-sided Mann–Whitney test (n = 31, Con; n = 31, *Mecp2* cKO; *P* = 0.8339). All data are presented as mean ± SEM. Individual points represent individual cells or sections.

To assess excitatory synaptic organization, we quantified immunohistochemical labeling of glutamatergic postsynaptic and presynaptic sites in the PC layer (PCL) and molecular cell layer (MCL). Our previous data indicated that the AMPA receptor subunits GluA1 and GluA4 are mainly expressed in Bergmann glia, whereas GluA2 is mainly expressed in PCs^32^. Consistent with the larger dendritic spines, GluA2 immunoreactivity was higher in *Mecp2* cKO mice (Fig. 7j). GluA4 was unchanged, whereas GluA1 was also higher, suggesting a potential non-cell-autonomous consequence of PC-specific *Mecp2* deletion on Bergmann glia (Fig. 7i,k). In contrast, the immunoreactivity of the vesicular glutamate transporters vGluT1, which labels parallel fiber (PF) presynaptic terminals, and vGluT2, which labels climbing fiber (CF) presynaptic terminals, did not differ between genotypes, and the relative extent of CF-associated vGluT2 labeling along PC dendrites was unchanged, arguing against a major reorganization of gross excitatory afferent patterning (Fig. 7l-n). Consistently, independent quantifications of the numerical density of vGluT1 and vGluT2 puncta did not show significant differences between genotypes (Extended Data Fig. 7a,b). These findings indicate that PC-specific *Mecp2* deletion causes selective structural and synaptic remodeling, pointing to altered postsynaptic integration and plasticity as a potential contributor to the observed deficits in associative learning.

### Altered intrinsic excitability of PCs in *Mecp2* cKO mice

To determine whether the structural remodeling of *Mecp2*-deficient PCs was associated with altered intrinsic properties, we performed intracellular whole-cell and extracellular cell-attached recordings in acute cerebellar slices. In response to hyperpolarizing and depolarizing current injections, *Mecp2* cKO PCs showed altered subthreshold voltage responses and firing output relative to controls (Fig. 8a,b). In particular, the rheobase was significantly smaller and the resting membrane potential was depolarized in *Mecp2* cKO PCs, whereas the threshold of Na^+^-dependent spikes was unchanged (Fig. 8c-e). These findings indicate that *Mecp2*-deficient PCs require less input to reach action potential discharge and are intrinsically more excitable at resting membrane potential.

**Fig. 8.**
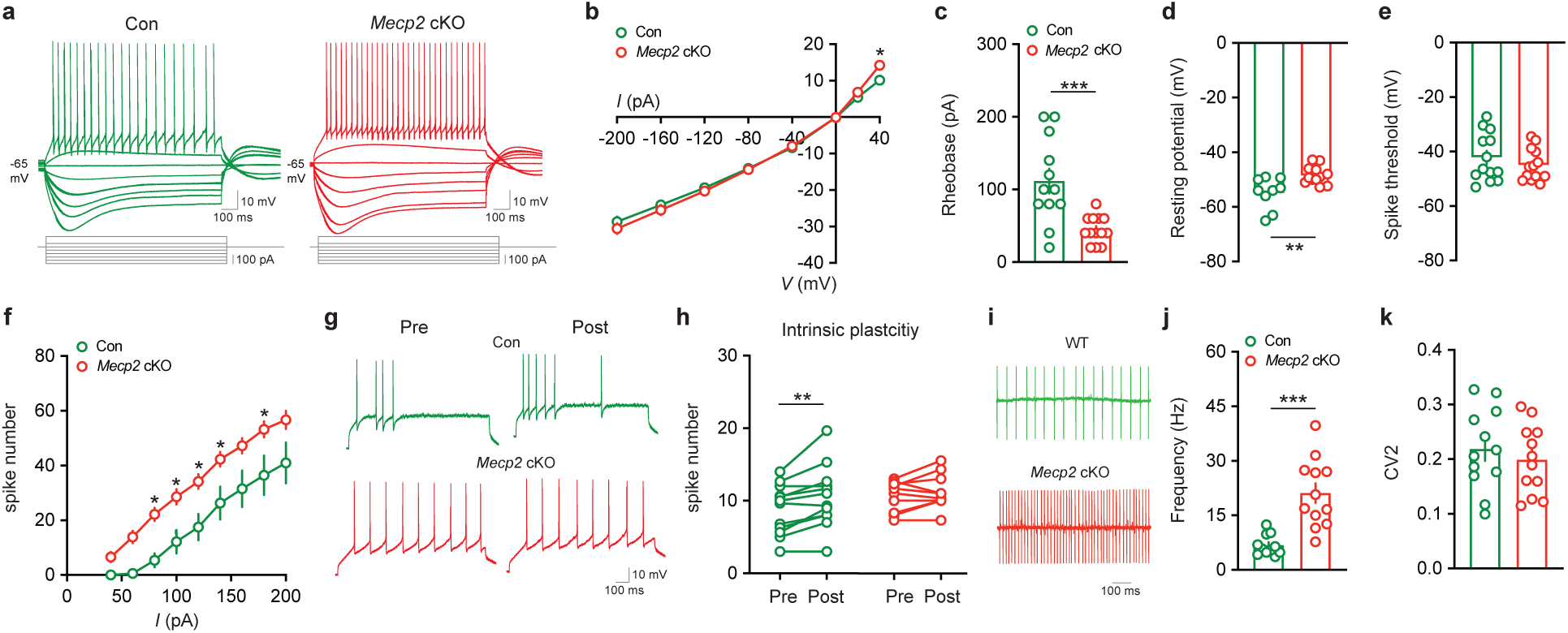
| Altered intrinsic excitability of PCs in *Mecp2* cKO mice. **a**, Representative subthreshold voltage responses and firing patterns evoked by current injections from – 200 to 100 pA. **b,** Input-output relationship between injected current and membrane voltage response. Two-way repeated-measures ANOVA with Bonferroni’s multiple-comparisons test revealed a significant current × genotype interaction *F*(7, 189) = 2.17, *P* = 0.0390, a significant main effect of input current *F*(7, 189) = 540.74, *P* < 0.001, but no main effect of genotype *F*(1, 17) = 0.06, *P* = 0.8137 (n = 13, Con; n = 14, *Mecp2* cKO). Post hoc testing identified a significant difference at 40 pA (*P* = 0.0442). **c,** Quantification of rheobase. Two-sided Mann-Whitney test (n = 12, Con; n = 13, *Mecp2* cKO; *P* < 0.001). **d,** Quantification of resting membrane potential. Unpaired two-sided Student’s *t*-test (n = 13, Con; n = 13, *Mecp2* cKO; *P* = 0.0035). **e,** Quantification of spike threshold. Unpaired two-sided Student’s *t*-test (n = 13, Con; n = 13, *Mecp2* cKO; *P* = 0.3563). **f,** Input-output relationship between injected current and action potential number (40-200 pA, 20 pA increments). Two-way repeated-measures ANOVA with Bonferroni’s multiple-comparisons test showed no significant current × genotype interaction *F*(8, 184) = 0.9165, *P* = 0.5040, but significant main effects of input current *F*(8, 184) = 92.09, *P* < 0.001 and genotype *F*(1, 23) = 10.07, *P* = 0.0042 (n = 12, Con; n = 13, *Mecp2* cKO). Post hoc testing identified significant differences at 80, 100, 120, 140, and 180 pA (*P* < 0.05). **g,** Representative test spikes recorded before and after the intrinsic plasticity induction protocol. **h,** Quantification of spike number before and after intrinsic plasticity induction. Paired two-sided Student’s *t*-test (n = 12, Con, *P* = 0.0048, Pre vs. Post; n = 13, *Mecp2* cKO, *P* = 0.1636, Pre vs. Post). **i,** Representative spontaneous spiking recorded in cell-attached mode. **j,** Quantification of spontaneous spike frequency. Two-sided Mann-Whitney test (n = 11, Con; n = 12, *Mecp2* cKO; *P* < 0.001). **k,** Quantification of CV2 of spontaneous spiking. Unpaired two-sided Student’s *t*-test (n = 12, Con; n = 12, *Mecp2* cKO; *P* = 0.4948). All data are presented as mean ± SEM. Individual points represent individual recordings.

Consistent with these data, current-evoked spike output was larger in *Mecp2* cKO PCs across a range of depolarizing steps (Fig. 8f). Additional passive membrane properties, as well as action potential waveform features, including spike amplitude, spike half-width, fast and slow afterhyperpolarization amplitudes, delay to first spike, and accommodation ratio, were all unchanged (Extended Data Fig. 8a-g). The amplitude of the voltage sag evoked by hyperpolarizing current injections was altered as a function of current step amplitude, further supporting a change in subthreshold membrane properties rather than a broad disruption of action potential generation or its waveform (Extended Data Fig. 8h). Thus, PC-specific *Mecp2* deletion selectively shifts intrinsic excitability without substantially altering the basic kinetics of Na^+^-dependent action potentials.

We next asked whether activity-dependent plasticity of intrinsic excitability was affected. In control PCs, repetitive depolarization increased subsequent spike output, representing typical intrinsic plasticity (Fig. 8g,h). In *Mecp2* cKO PCs, however, the same induction protocol failed to enhance firing, indicating that *Mecp2* loss impairs the ability of PCs to undergo activity-dependent increases in intrinsic excitability.

This altered excitability profile was also evident for spontaneously occurring Na^+^-dependent action potentials. Cell-attached extracellular recordings showed that *Mecp2* cKO PCs fired spontaneously at a substantially higher frequency than controls, whereas spike train variability, measured by their coefficient of variance (CV^2^), was unchanged (Fig. 8i-k). Thus, *Mecp2*-deficient PCs are not only hyperexcitable in response to injected current but also exhibit elevated spontaneous firing without a major change in firing regularity. Together, these findings indicate that PC-specific *Mecp2* deletion causes a selective increase in intrinsic excitability while impairing intrinsic plasticity, consistent with their structural remodeling.

### Impaired synaptic transmission and plasticity in *Mecp2* cKO Purkinje cells

To determine whether synaptic function was altered in *Mecp2* cKO PCs, we examined PF– and CF-mediated transmission onto PCs by intracellular whole-cell recordings in *ex vivo* cerebellar slices. Analysis of PF-PC input-output relationships showed that the amplitude of excitatory postsynaptic currents (EPSC) evoked by PF stimulation was larger in *Mecp2* cKO PCs, particularly at higher stimulus intensities (Fig. 9a), indicating atypically stronger PF-PC synapses. The ratio of EPSCs evoked by pairs of PF stimulation was unchanged across all inter-pulse intervals tested (Fig. 9b), consistent with the unaltered levels of presynaptic vGluT1 expression, which argues against major consequences for presynaptic function at PF-PC synapses and suggests higher postsynaptic sensitivity.

**Fig. 9.**
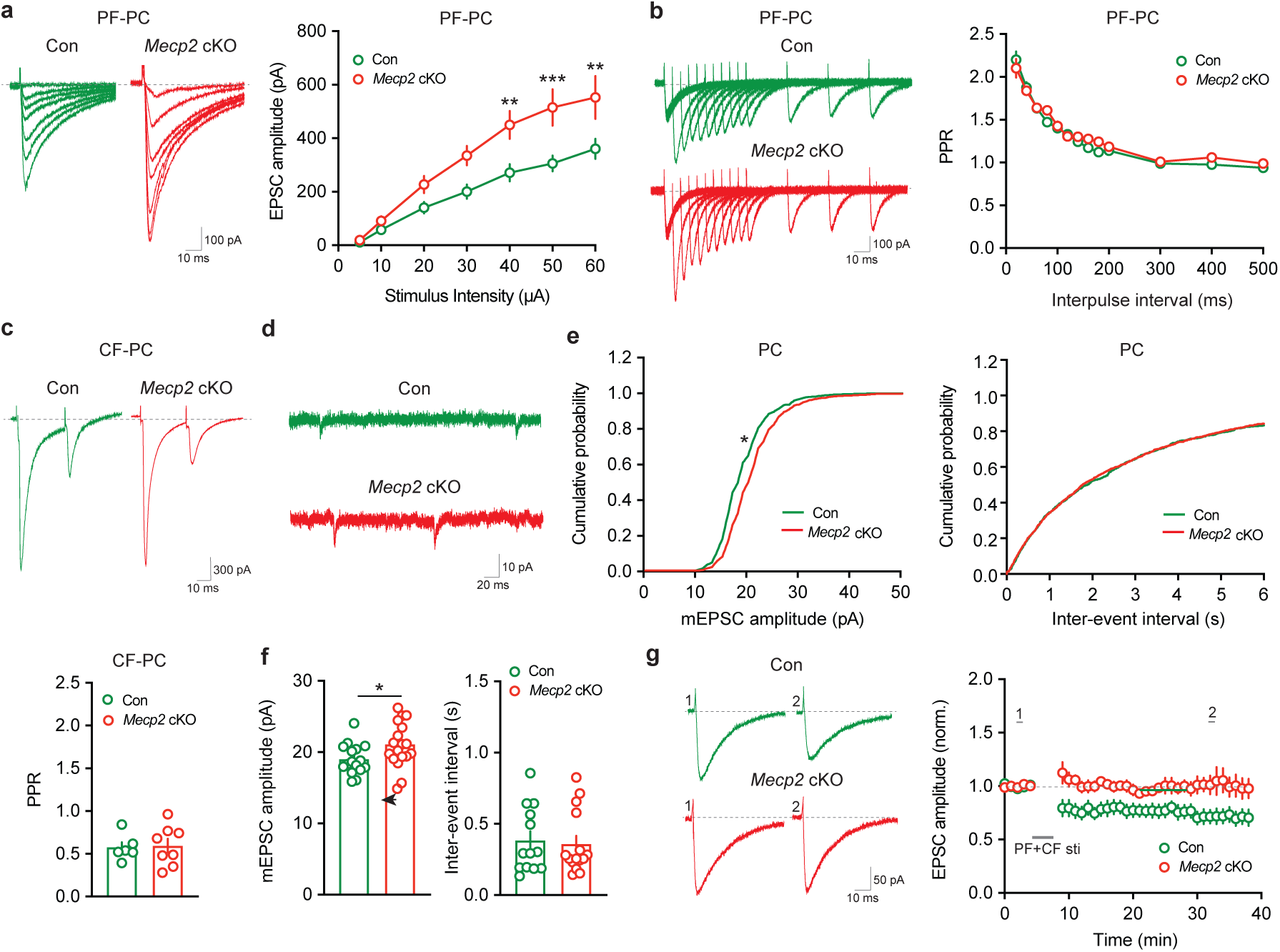
| PF and CF synaptic properties in PCs of *Mecp2* cKO mice. **a**, Representative PF-PC EPSCs evoked by increasing stimulus intensities and quantification of the PF-PC input-output relationship. Two-way repeated-measures ANOVA with Bonferroni’s multiple-comparisons test showed a significant stimulus intensity × genotype interaction *F*(6, 293) = 2.78, *P* = 0.0120, a significant main effect of stimulus intensity *F*(6, 293) = 44.31, *P* < 0.001, and a significant main effect of genotype *F*(1, 293) = 43.88, *P* < 0.001 (n = 35, Con; n = 12, *Mecp2* cKO). Post hoc testing identified significant differences at 40, 50, and 60 μA (*P* < 0.05). **b,** Representative paired-pulse PF-PC EPSCs and quantification of paired-pulse ratio (PPR) across inter-pulse intervals. Two-way repeated-measures ANOVA with Bonferroni’s multiple-comparisons test showed no significant inter-pulse interval × genotype interaction *F*(17, 353) = 1.140, *P* = 0.3133, a significant main effect of inter-pulse interval *F*(17, 353) = 124.7, *P* < 0.001, and no significant main effect of genotype *F*(1, 25) = 1.071, *P* = 0.3106 (n = 15, Con; n = 12, *Mecp2* cKO). **c,** Representative paired-pulse CF-PC EPSCs and quantification of CF paired-pulse ratio at a 40-ms inter-pulse interval. Unpaired two-sided Student’s *t*-test (n = 6, Con; n = 8, *Mecp2* cKO; *P* = 0.8642). **d,** Representative mEPSC traces recorded from PCs. **e,** Cumulative probability plots of mEPSC amplitude and inter-event interval. Kolmogorov-Smirnov test (n = 11, Con; n = 16, *Mecp2* cKO; *P* = 0.0124, mEPSC amplitude; *P* = 0.4009, inter-event interval). **f,** Quantification of mean mEPSC amplitude and inter-event interval. Unpaired two-sided Student’s *t*-test (n = 16, Con; n = 18, *Mecp2* cKO; *P* = 0.0496, mEPSC amplitude; *P* = 0.7685, inter-event interval). **g,** Representative PF-LTD traces and summary time course of normalized PF-PC EPSC amplitude before and after LTD induction. Two-way repeated-measures ANOVA showed a significant time ξ genotype interaction *F*(34, 612) = 2.570, *P* < 0.001, a significant main effect of time *F*(34, 612) = 2.493, *P* < 0.001, and a significant main effect of genotype *F*(1, 18) = 9.329, *P* < 0.0068 (n = 12, Con; n = 8, *Mecp2* cKO). All data are presented as mean ± SEM. Individual points represent individual recordings.

We next assessed the properties of CF-PC synapses. The threshold stimulus intensity required to evoke EPSCs in PCs by CF stimulation was unchanged (Extended Data Fig. 9a,b), as was the proportion of PCs showing one-step (i.e. all-or-none) responses versus graded responses reflecting multi-innervation of individual PCs by CFs (Extended Data Fig. 9c). Consistent with preserved vGluT2 expression, the paired-pulse ratio of CF-evoked EPSCs was also unaffected (Fig. 9c). These findings indicate that gross CF innervation pattern and short-term CF synaptic plasticity remain largely intact following PC-specific *Mecp2* deletion.

To further characterize excitatory synaptic drive onto PCs, we recorded spontaneous miniature EPSCs originating from both PFs and CFs in the absence of Na^+^-dependent action potentials (Fig. 9d). mEPSC amplitudes were larger in *Mecp2* cKO PCs, whereas their inter-event intervals were unchanged (Fig. 9e,f), again supporting enhanced postsynaptic responsiveness without detectable changes in presynaptic release probability. This pattern is consistent with the structural findings of larger spines and higher GluA2 immunoreactivity in *Mecp2* cKO PCs.

We then determined whether long-term plasticity at PF-PC synapses was altered. Paired PF-CF stimulation for 5 min at 1 Hz allowing PCs to fire action potentials evokes the expected long-term depression (LTD) of EPSC amplitude in control mice, but *Mecp2* cKO PCs failed to show this typical LTD (Fig. 9g). Thus, *Mecp2* deficiency in PCs also disrupts long-term plasticity at PF-PC synapses. These results are consistent with the morphological and intrinsic property consequences observed in *Mecp2* cKO PCs and suggest that atypical postsynaptic integration and impaired synaptic plasticity in cerebellar PCs contribute to the observed motor learning deficits in these mice.

## Discussion

Our study identifies PC-selective *Mecp2* loss as a multiscale driver of cerebellar dysfunction in RTT-related motor phenotypes. Across behavior, circuit activity, and cellular physiology, the phenotype was selective rather than global. PC-specific *Mecp2* deletion spared gross locomotor activity, anxiety-like behavior, sociability, and gross motor strength. In contrast, it disrupted higher-order gait coordination, impaired rotarod learning, delayed delay eyeblink conditioning, and produced a more persistent deficit in trace eyeblink conditioning. At the circuit level, these behavioral consequences were accompanied by a failure of learning-related reorganization of CxSpk signaling, and by blunted CS-related population Ca^2+^ dynamics during associative learning. At the cellular level, *Mecp2*-deficient PCs showed lower dendritic complexity, larger dendritic spines, higher glutamatergic receptor levels, enhanced intrinsic excitability, impaired plasticity of intrinsic properties, stronger PF drive, and impaired PF-PC long-term synaptic plasticity. These findings suggest that *Mecp2* loss in PCs does not simply weaken cerebellar function in a nonspecific manner but instead alters the rules by which cerebellar output neurons encode error, integrate synaptic input, and support adaptive motor learning.

A notable feature of the behavioral phenotype in mice with selective deletion of *Mecp2* in PCs is that it emerges first as a subtle coordination defect rather than as a gross motor syndrome. Standard assays failed to reveal broad locomotor abnormalities, but *CatWalk* analysis uncovered selective deficits in stance-phase control, interlimb timing, paw placement and support configuration, while leaving stride length, step cycle and overall regularity relatively preserved. This pattern is consistent with a cerebellar deficit in fine movement calibration and predictive coordination rather than in movement generation itself^18, 33^. Such a phenotype fits the canonical role of the cerebellum in timing, coordination, and adaptive control, and is also consistent with the clinical observation that RTT motor dysfunction often includes impaired gait precision, ataxia, and deterioration of learned movement before complete loss of gross movement capacity^34–36^. The motor learning phenotype was similarly selective. Skilled single-pellet reaching was largely preserved, with intact reach distance and movement duration, and only a modest overall reduction in success rate, whereas rotarod learning was clearly impaired and eyeblink conditioning was even more vulnerable. This task hierarchy is mechanistically informative. Skilled reaching depends heavily on corticospinal and motor cortical circuits, with cerebellar contributions biased toward movement refinement and online correction^26, 27^, whereas rotarod and eyeblink conditioning place stronger demands on cerebellar timing, adaptation and plasticity^28, 37^. The finding that trace eyeblink conditioning was more severely affected than delay conditioning suggests that *Mecp2* loss in PCs becomes especially consequential when cerebellar computations must be integrated with forebrain networks that bridge temporally separated events^29, 38^. Thus, the phenotype is not simply “motor” vs. “non-motor” behavior, or delay vs. trace conditioning; instead, it is strongest in behaviors that require precise cerebellar teaching signals coupled to temporally extended associative processing.

These observations help reconcile our findings with two recent studies of the consequences of *Mecp2* deletion in the cerebellum^23, 24^. Achilly et al. found that non-motor social behaviors remained intact after PC-related cerebellar *Mecp2* manipulation, which is consistent with our results^23^. However, they showed that deleting *Mecp2* from the cerebellum as a whole impairs rotarod and delay eyeblink motor learning, whereas deletion from individual cerebellar neuronal subtypes, including PCs, did not affect rotarod learning. Whether PC-specific *Mecp2* deletion affects delay or trace eyeblink conditioning was not addressed in that study. By contrast, Xu et al. reported that PC-specific *Mecp2* deletion is sufficient to impair motor learning and produce social deficits^24^. Our results agree with Xu et al. in showing that PC-selective *Mecp2* loss is sufficient to impair motor learning but differ from Xu et al. regarding the social behavior phenotype. These differences may reflect, at least in part, the ages at which the experiments were performed, because the behavioral consequences of *Mecp2* deletion may evolve during development and may also be shaped by compensatory processes. Achilly et al. primarily examined mice at around 6 months of age, our experiments were performed at 3-5 months, and Xu et al. studied younger mice, mostly under 3 months. Consistent with these behavioral differences, some PC cellular and intrinsic properties also differ across studies, which may additionally reflect differences in assay sensitivity, cerebellar region, or physiological readout. Our study also extends these prior reports in two important ways. First, we show that the phenotype is not uniform across behavioral domains: pellet-reaching is largely spared, rotarod and delay eyeblink conditioning are impaired, and trace conditioning is most severely affected. Second, the consequences are not limited to intrinsic excitability; it also includes disruption of learning-related CxSpk reorganization, atypical population-level Ca^2+^ dynamics, structural remodeling, and impaired long-term synaptic plasticity.

The eyeblink data provide the clearest mechanistic link between altered PC physiology and impaired learning. In control mice, CxSpk activity shifted over training from a predominantly US-aligned response toward an enhanced CS-aligned response. This redistribution is consistent with previous work showing that, after conditioning, CFs can acquire delayed CS-evoked responses while their probability of responding to the US is reduced, suggesting that CxSpks may carry learned predictive or timing-related teaching signals^39–42^. In *Mecp2* cKO PCs, however, this forward shift was impaired. Because CF-triggered CxSpks are central to the induction of PF-PC LTD in many models of eyeblink conditioning^43–46^, failure to appropriately re-time this signal would be expected to weaken associative plasticity at PF-PC synapses and thereby impair conditioned response (CR) acquisition. Thus, the behavioral, *in vivo* physiological, and *ex vivo* synaptic findings converge on a common mechanism in which *Mecp2* loss in PCs weakens the conversion of CF-related instructive signals into stable learning-related synaptic change. At the same time, current models of cerebellar function indicate that this shift in CxSpk activity can also contribute to conditioned SmSpk suppression^39, 43, 47, 48^. However, in our dataset, the increase in CS-related CxSpks after learning was not accompanied by a robust overall decrease in SmSpk firing. There are at least two possible explanations for this. First, only a subset of CxSpk appears to shift forward into the CS epoch, so their net effect on mean SmSpk output may be diluted when activity is averaged across all trials. Indeed, the expected CxSpk-induced suppression becomes clear only when analysis is restricted to the subset of trials in which the relevant CxSpk occurs. Second, SmSpk firing during behavior is shaped not only by CF-triggered pauses, but also by CF drive, eyelid movement, body movement, locomotor state and other sensorimotor variables^20, 21, 49^. Consistent with this interpretation, even when US-related CxSpk activity remains prominent, SmSpk firing does not necessarily decrease overall. Our data further show that running speed changes after both the CS and US with training, which could additionally influence SmSpk output. Thus, the absence of gross population SmSpk suppression does not argue against learning-related CxSpk reorganization; rather, it suggests that this effect is embedded within a richer behavioral state space and may be masked by concurrent excitatory and movement-related influences.

Our data also show that neither single-cell SmSpk nor CxSpk responses correlated strongly with CR percentage or running speed, whereas the population Ca^2+^ signal did. The absence of strong single-cell correlations likely reflects the fact that any individual PC captures only a limited component of the ensemble code, whereas behavior emerges from coordinated activity across a broader microzonal population. The significant relationship between CS-aligned Ca^2+^ responses and CR percentage, together with the acquisition-related increase in CS signals and decrease in US signals, suggests that the relevant learning signal is represented primarily at the level of a coordinated PC ensemble rather than at the level of any single neuron. Thus, as learning proceeds, CxSpks probability may increase across a neuronal ensemble during the CS epoch even though CR percentage is not linearly predictable from the firing of an individual recorded cell^42^. Population Ca^2+^ photometry recordings revealed two additional features that are conceptually important. First, consistent with the single-unit recordings, the learning-related increase in CS-aligned activity was blunted in *Mecp2* cKO mice. An unexpected observation, however, was that US-related population Ca^2+^ signals declined modestly across training in both genotypes. This differs from the single-unit results, in which the clearest learning-related reorganization of CxSpk signaling was observed in controls and was largely absent in cKOs. One possible explanation is that GCaMP photometry captures a broader mixture of signals, including CxSpk-related dendritic Ca^2+^ elevations, subthreshold dendritic events, and wider state-dependent or network-driven activity, whereas extracellular single-unit recording isolates the spiking output of individual PCs^50,51^. In this context, the residual learning-related Ca^2+^ photometry signal in cKO mice may reflect preserved but weaker ensemble recruitment, dendritic signaling that is not efficiently converted into CxSpk output, or population averaging across multiple microzones. Second, our bilateral recordings showed that learning-related CS– and US-related signals were strongest ipsilateral to the air puff but were also detectable contralaterally, albeit to a lesser extent. This pattern likely reflects the fact that eyeblink conditioning engages distributed cerebellar networks rather than a strictly unilateral local circuit. The stronger genotype effect on the ipsilateral side therefore fits a lateralized teaching signal superimposed on a broader bilateral state representation.

At the cellular level, the phenotype points strongly to a postsynaptic consequence within PCs. Morphologically, *Mecp2* cKO PCs exhibited simpler dendritic complexity and smaller coverage, larger dendritic spines, and higher GluA2 immunoreactivity, with a more modest increase in GluA1. Functionally, *Mecp2*-lacking PCs showed lower rheobase, a depolarized resting membrane potential, increased current-evoked firing, higher spontaneous firing, and impaired plasticity of intrinsic properties. At the synaptic level, PF-PC synapses were stronger, with steeper input-output curves and larger mEPSCs, whereas short-term plasticity (paired-pulse ratios) at both PF-PC and CF-PC synapses, vGluT1 and vGluT2 immunoreactivity, puncta density, and the gross CF innervation pattern were all preserved. On the other hand, PF-LTD was impaired in *Mecp2*-lacking PCs. This overall profile argues that the dominant effect of PC-specific *Mecp2* deletion is not a presynaptic remodeling of PF or CF afferents, but rather a shift in postsynaptic integration and excitability within PCs themselves. These postsynaptic changes provide a plausible substrate for the deficits in synaptic plasticity. In a PC that is already hyperexcitable and receiving stronger PF-driven depolarization, the balance of signaling mechanisms that normally supports bidirectional synaptic plasticity is likely shifted, producing a maladaptive change in the plasticity landscape^52^. This interpretation is consistent with our previous findings in the hippocampus of *Mecp2* knockout mice^53, 54^. The GluA1 result further raises the possibility of a non-cell-autonomous component to the phenotype, because our prior work indicates that GluA2 is enriched in PCs, whereas GluA1 and GluA4 are enriched in Bergmann glia^32^. This distinction is important because it suggests that, although the genetic manipulation is PC specific, part of the resulting cerebellar phenotype may emerge from altered communication within the PC-Bergmann glia network. Thus, *Mecp2* loss in PCs may have both cell-autonomous consequences for excitability and synaptic integration, as well as secondary non-cell-autonomous consequences for local glial signaling.

Overall, our findings support a model in which PC-specific *Mecp2* loss preserves general motor capacity but destabilizes the adaptive computations required for cerebellar learning. Structural remodeling increased postsynaptic responsiveness, and elevated intrinsic excitability shift PCs into a high-gain state, whereas impaired intrinsic plasticity, impaired PF-PC LTD, and disrupted learning-related CxSpk reorganization limit their ability to update output during training. In this framework, *Mecp2*-deficient PCs can still support basic movement and partially learned behaviors, but they fail to properly transform instructive signals into stable changes in cerebellar network activity. The resulting phenotype is therefore not a complete motor failure, but reduced precision, poorer coordination, and impaired adaptive updating, especially when behaviors require temporally bridged associations or distributed cerebello-forebrain interactions. These results place PCs at the center of RTT-related cerebellar dysfunction and suggest that RTT motor phenotypes arise not only from cortical or basal ganglia pathology, but also from disrupted computation within cerebellar output neurons.

## Methods

### Animal models

All mice were handled and housed in accordance with National Institutes of Health guidelines. Both male and female mice were used in this study at the age of 3-5 months. Mice were maintained at 25 °C under a 12-h light/dark cycle with 40-60% humidity and provided ad libitum access to food and water. All procedures were approved by the Institutional Animal Care and Use Committees of the University of Alabama at Birmingham (IACUC-22247) and Michigan State University (PROTO202500345). The generation and genotyping of floxed *Mecp2* mice (B6;129P2*-Mecp2^tm1Bird^*/J), *Pcp2*-Cre mice, and Ai95D (B6J.Cg-*Gt(ROSA)26Sor^tm95.1(CAG-GCaMP6f)Hze/MwarJ)^* have been described previously^8, 25, 55^. Genotyping and husbandry were performed according to vendor protocols. Heterozygous *Pcp2*-Cre offspring were used for breeding and experiments. To generate PC-specific *Mecp2* conditional knockout (cKO) mice, *Pcp2*-Cre mice were crossed with *Mecp2* floxed mice. Female homozygous *Pcp2*-Cre; *Mecp2*^fl/fl^ mice and male hemizygous *Pcp2*-Cre; *Mecp2*^fl/y^ mice were used as cKO animals, along with littermate controls. For Ca^2+^ photometry experiments, offspring were further crossed with homozygous Ai95D mice to generate triple-transgenic animals expressing GCaMP6f in PCs.

### Open field test

Mice were placed in the center of a 30 × 40 cm open-field arena. Following a 10-min habituation period, behavior was recorded for 10 min using an infrared-sensitive Gigabit Ethernet video camera (ace acA780-75gm, Basler). The arena floor was virtually divided into nine zones. Behavioral measures included total distance traveled, mean velocity, immobility time, time spent in the center zone, number of center entries, and distance traveled in the center zone normalized to total distance traveled. Center time was normalized to total recording time. Rearing and grooming were manually scored, and their frequency and duration were analyzed as indicated. Tracking and analyses were performed using EthoVision XT 16 software (Noldus).

### Three-chamber social test

Mice were acclimated to handling for 3 consecutive days before testing. Social behavior was assessed in a three-chamber apparatus (60 × 30 × 20 cm) consisting of a center chamber and two side chambers connected by removable partitions. At the start of the assay, each mouse was placed in the center chamber and allowed to freely explore the entire apparatus for 5 min for habituation. For the sociability phase, the test mouse was gently guided back to the center chamber and temporarily confined there while a novel stimulus mouse was placed under a wire pencil cup in one side chamber; an identical empty cup was placed in the opposite side chamber. The partitions were then removed, and the test mouse was allowed to freely explore all three chambers for 10 min. After this session, the test mouse was again returned to the center chamber and briefly confined. For the social memory phase, a second novel stimulus mouse was placed under the previously empty cup, while the first novel mouse remained in its original location. The partitions were removed, and the test mouse was allowed to explore the apparatus for an additional 10 min.

Behavior was recorded and analyzed using EthoVision XT 16 software. Social investigation was quantified as the time spent sniffing each cup. Sociability was assessed by comparing investigation time directed toward the cup containing the first novel mouse versus the empty cup. Social memory was assessed by comparing investigation time directed toward the second novel mouse versus the now-familiar first novel mouse. Preference indices were calculated as follows: for sociability,

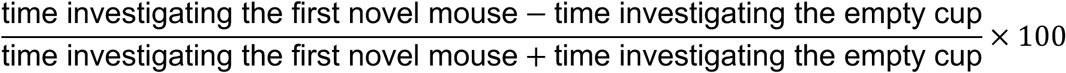

and for social memory,

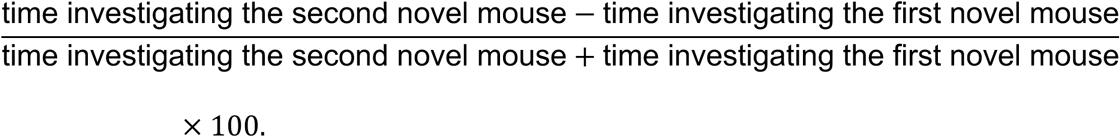

### Grip strength test

Forelimb grip strength was assessed using a grip strength meter (San Diego Instruments). Mice were allowed to grasp the grid of the apparatus with their forepaws and were then gently pulled backward by the tail until they released the grid. Peak force was recorded for each trial. Each mouse was tested in five trials, and the average peak force was used as the measure of forelimb strength.

### Ladder rung walking test

To assess skilled walking and limb placement, mice were tested on a horizontal ladder rung apparatus (Bio-FMA, Bioseb). Each mouse was placed at one end of the ladder and allowed to walk freely to the opposite end. During traversal, movement and paw placement errors were automatically detected by infrared sensor arrays positioned above and below the ladder. Mice were given three trials separated by 10-min inter-trial intervals. Foot-slip errors were quantified separately for the forelimbs and hindlimbs, and the mean number of errors across trials was used as the measure of motor coordination and stepping accuracy.

### CatWalk gait analysis

Gait performance was assessed using an automated CatWalk XT system (Noldus)^56^. The system consists of an enclosed, internally illuminated glass walkway, where paw contacts scatter light and are captured by a high-speed camera positioned beneath the platform. All behavioral testing was conducted in a quiet, low-light environment. Mice were habituated to the apparatus and trained to traverse the walkway for 3 consecutive days. On the test day, each mouse completed three compliant runs, defined as uninterrupted crossings without stopping, turning, or hesitating and within a consistent speed range. Trials were separated by 10-min inter-trial intervals. Only compliant runs were included, and all parameters were averaged across runs for each animal.

Paw contacts were automatically detected and classified as left forepaw (LF), left hind paw (LH), right forepaw (RF), and right hind paw (RH) using CatWalk XT software, followed by manual verification of paw labeling. Gait parameters were automatically extracted and grouped into four categories:

(1) Run characteristics and kinetic parameters included stride length, print area, print position, and swing speed. Stride length was defined as the distance between consecutive placements of the same paw. Print area reflected the surface area of paw contact, and swing speed was calculated during the swing phase.
(2) Temporal parameters included stand time, swing time, step cycle, and maximal contact relative to stand time. The step cycle was defined as the interval between successive contacts of the same paw and consisted of stand (paw contact) and swing (no contact) phases. Maximal contact was defined as the time point of peak paw contact intensity during the stance phase, normalized to stand time.
(3) Spatial parameters included base of support (BOS) and print position. BOS was calculated separately for forepaws and hind paws as the average distance between left and right paws. Print position was defined as the relative placement of the hind paw with respect to the preceding forepaw.
(4) Interlimb coordination parameters included phase dispersion, support patterns, and regularity index. Phase dispersion quantified the temporal relationship between limb pairs (LF→LH, LF→RF, LF→RH, LH→RH, RF→LH, RF→RH) as a percentage of the step cycle. Support patterns represented the proportion of time spent in different paw-support configurations (zero-, single-, diagonal-, girdle-, lateral-, three-, and four-paw support). The regularity index was used as a measure of step sequence consistency.

Step sequence patterns (Aa, Ab, Ca, Cb, Ra, and Rb) were additionally quantified to characterize locomotor coordination. These were defined as follows: Aa (LF–LH–RF–RH), Ab (LF–RH–RF–LH), Ca (RF–LF–RH–LH), Cb (LF–RF–LH–RH), Ra (RF–LF–LH–RH), and Rb (LF–RF–RH–LH).

### Single-pellet reaching task and kinematic analysis

Mice were housed in small groups (3-4 per cage) and handled daily for at least 2 days before behavioral testing. To increase motivation for food retrieval, mice were maintained under controlled food restriction using the task food reward at ∼80% of their baseline body weight throughout the experiment, and body weight was monitored daily. Training consisted of three phases: habituation, shaping, and task acquisition. During habituation, mice were placed individually in the reaching chamber under dim lighting each day and allowed to consume up to 20 food pellets scattered on the chamber floor for 30 min per session. Habituation continued until mice reliably consumed ≥75% of the pellets. During shaping, forelimb preference was determined by presenting pellets just outside the reaching slot and recording the paw used for repeated reaching attempts. The dominant forelimb was defined as the paw used in the majority of attempts, and the pellet delivery position was then aligned contralateral to the preferred paw to enforce unilateral reaching. Mice were subsequently trained on an automated single-pellet reaching task. Pellet delivery was triggered when the mouse moved to the rear of the chamber and interrupted an infrared beam, causing a pellet to be raised into position at the reaching slot. Mice were trained for 20 min per day over 12 days to reach, grasp, and retrieve pellets using the preferred forelimb. Reaching outcomes were categorized as success (grasp and retrieval), drop (grasp without retrieval), fail (miss or displacement of the pellet), or no attempt.

Forelimb movements were recorded using a dual-camera system positioned above and lateral to the reaching chamber, enabling three-dimensional reconstruction of reaching trajectories. Videos were acquired at 150 frames per second and segmented into individual trials for analysis. Forelimb kinematics were extracted using DeepLabCut, which tracked the positions of individual digits, the distal interphalangeal joint of the second digit, the forepaw, and the pellet in each frame. Separate models for the top-down and lateral camera views were developed and iteratively refined until stable tracking performance was achieved across multiple animals and sessions. Extracted coordinates from both camera views were then combined using custom MATLAB scripts to reconstruct 3D trajectories^57, 58^.

To ensure data quality during trajectory processing, labeled points with low confidence scores (< 0.80) were filtered to exclude low-confidence detections. Such points were interpreted as instances in which the tracked feature was out of frame or the pellet was not present for a valid reaching attempt. The filtered preliminary data were cleaned using linear interpolation (filloutliers function, MATLAB) and then analyzed to identify the start and end points of reaching movements within each trial. Trials were considered valid if the pellet appeared within the expected time window at the beginning of the trial and remained in position until a reach attempt began. Only the first reaching attempt in each trial with food present was analyzed. Filtered trajectories were then smoothed using a Savitzky–Golay filter, with a moving average filter applied when necessary to remove noise and outliers.

Reach and grasp phases of movement were differentiated using custom angle-based kinematics^57^. For valid trials, the angle formed by the forepaw, distal interphalangeal joint of the second digit, and the digit tip was calculated for each frame. Reaching onset was defined as the initial frame in which the paw crossed the reaching slot into the task space. The reaching phase ended at the frame corresponding to the maximal local joint angle, which represented peak extension and the transition from reach to grasp. The subsequent local minimum angle was used to define the end of grasp. Three-dimensional forelimb trajectories were then interpolated to 100 evenly spaced points using piecewise cubic Hermite interpolation (pchip) to standardize trajectory length across trials. The following kinematic parameters were extracted: (1) reach time, defined as the duration from reach onset to peak extension; (2) grasp time, defined as the duration from peak extension to grasp completion; (3) reach arc, defined as the total distance traveled during the reaching phase; (4) grasp arc, defined as the total distance traveled during the grasping phase; (5) total trajectory length, defined as the combined distance traveled during the reaching and grasping phases; and (6) movement variability, quantified as the average deviation of individual trajectories from the mean trajectory across trials and calculated as the average absolute distance between each trajectory and the mean trajectory over time. Trial-level measurements were averaged within each session to obtain daily performance metrics, including mean reach distance, grasp distance, reach time, and grasp time, and were then averaged across animals for group comparisons. Three-dimensional trajectories were further processed in Python with cubic spline interpolation for visualization of smoothed trajectories.

### Rotarod test

Mice were trained on a rotarod apparatus (ROTOR-ROD, San Diego Instruments) for 3 consecutive days. During each trial, mice were placed on the rotating rod for up to 1 min and returned to their home cage between trials. Mice underwent 4 trials per day with a 10-min inter-trial interval. On the following day, mice were subjected to 3 test trials. Latency to fall was recorded for each trial, and the average latency across trials was used for analysis.

### Delay and trace eyeblink conditioning

Stereotactic surgery was performed to implant a custom-built stainless steel headplate, as described previously^59^. Briefly, mice were anesthetized with isoflurane (3% for induction, 1% for maintenance), and secured in a stereotactic frame, and body temperature was maintained using a heating pad. Ophthalmic ointment was applied to prevent corneal drying. A midline incision was made to expose the skull, and the overlying fascia was removed. The headplate was secured on the skull using miniature screws and C&B Metabond dental cement (Parkell). The incision was then closed with surgical glue and sutures, and mice were allowed to recover for 7 days before behavioral training.

Following recovery, mice were acclimated to head restraint and allowed to run freely on a foam cylinder for 2 days. Eyeblink conditioning was then conducted in daily sessions under head-fixed conditions. Mice underwent a continuous training paradigm consisting of 12 acquisition sessions, 3 extinction sessions, and 2 reacquisition sessions. Each session consisted of 110 trials with a randomized inter-trial interval of 10-15 s. During acquisition and reacquisition, a conditioned stimulus (CS) was paired with an unconditioned stimulus (US) in each trial, except for every 10th trial, in which the CS was presented alone to assess CRs. During extinction sessions, the CS was presented without the US in all trials. For delay eyeblink conditioning, the CS consisted of a 350-ms blue LED light (M470F1, Thorlabs), and the US consisted of a 30-ms air puff (6-8 psi) delivered through a 23-gauge needle. The CS and US were co-terminated. For trace eyeblink conditioning, the CS consisted of a 50-ms light stimulus, followed by a 270-ms stimulus-free interval, and then a 30-ms air puff US.

Training procedure was performed under infrared illumination (IR56-56, CMVision). Eyelid movements were captured using a high-speed monochrome camera (250 frames/s; Mako G-040, Allied Vision) equipped with a 25-mm lens. Image acquisition was controlled using the MATLAB Image Acquisition Toolbox with custom-written scripts, which extracted subframes containing only the eye and surrounding region. Offline analysis was performed using custom MATLAB code. Each frame was converted to a binary image by applying a threshold that distinguished the eye region from the surrounding fur. Pixel values within the eye region were summed for each frame, and baseline values were calculated from the 100-ms period preceding CS onset. Eyelid closure was then expressed as the percentage change relative to baseline, such that 0% corresponded to the fully open state and 100% corresponded to complete eyelid closure. CRs were defined as trials in which eyelid closure exceeded 10% during the CS period before US onset. The percentage of CRs for each session was calculated as the number of CR-positive trials divided by the total number of trials. CR amplitude was quantified from CS-only trials (every 10th trial) as the peak eyelid closure (%) reached during the CS period.

### Acute *In vivo* single-unit recording during eyeblink conditioning

Mice were prepared for head-fixed *in vivo* recordings using the headplate implantation procedure described above, with additional preparation for acute electrophysiological recording^60^. Briefly, stainless steel skull screws were implanted bilaterally posterior to bregma, with one screw serving as a reference electrode. A silver wire was wrapped around the reference screw and secured with conductive epoxy to provide stable electrical grounding. A craniotomy (∼3 mm diameter) was made over the cerebellum (centered approximately 6.5 mm posterior and 2.0 mm lateral to bregma), targeting lobules involved in eyeblink conditioning. The skull bone was carefully removed while preserving the dura. A custom recording chamber was constructed around the craniotomy using dental acrylic, and the exposed dura was protected with sterile silicone elastomer (Kwik-Sil) between recording sessions.

Two *in vivo* recordings were performed during early (EA, days 1–2) and late (LA, days 9–12) acquisition phases of eyeblink conditioning, representing pre– and post-learning stages. Acute extracellular recordings were obtained using monopolar tungsten microelectrodes (2–5 MΩ impedance at 1 kHz; FHC). Electrodes were advanced into the cerebellum using a motorized micromanipulator (MP-285, Sutter Instrument) under visual guidance with a digital microscope (Dino-Lite). Penetrations were systematically spaced (∼100-200 μm) to sample across the target region. Electrodes were lowered gradually while monitoring neural activity to identify cerebellar layers and isolate single units. Neural signals were amplified using an AC/DC differential amplifier (Model 3000, A-M Systems), band-pass filtered, and digitized at 40 kHz for offline analysis. All components were grounded to a common reference to minimize electrical noise. PCs were identified based on established electrophysiological criteria, including (1) high-frequency SmSpk firing (∼50-100 Hz), (2) the presence of low-frequency CxSpks (∼1 Hz), and (3) a characteristic pause in SmSpks firing (10-30 ms) following each CxSpk. Only well-isolated units with stable waveform and firing properties throughout the recording session were included for analysis.

Stimulus delivery (CS and US), eyelid movement recordings, treadmill movement measured by a rotary encoder, and electrophysiological signals were synchronized using TTL pulses to ensure precise temporal alignment. Spike times were aligned to CS and US onset for analysis using custom MATLAB and Python scripts. For each recording, peri-event time histograms (PETHs) and smoothed firing rate profiles were computed separately for SmSpks and CxSpks. For CS– and US-aligned analyses, spike counts were calculated in 20-ms bins within defined time windows, including a 100-ms pre-CS baseline period and a 160-ms post-CS or post-US window during acquisition sessions with air-puff stimulation. Firing rates were calculated by normalizing spike counts to bin duration and number of trials and then normalized to baseline activity. Normalized CS– and US-aligned SmSpk and CxSpk responses were analyzed separately for early and late acquisition phases and compared with behavioral measures, including CR (%) and locomotor speed. Data were averaged across trials within each session and then across sessions within EA and LA phases for group-level analysis.

### *In vivo* fiber photometry during eyeblink conditioning

Mice expressing GCaMP6f in PCs underwent surgical procedures as described above. Fiber optic cannulas (200 μm core diameter, 0.37 NA; RWD Life Science) were implanted bilaterally over cerebellar lobule VI involved in eyeblink conditioning. Mice were allowed to recover for 1 week prior to behavioral training, and only animals exhibiting robust GCaMP6f fluorescence signals were included in subsequent experiments.

Fiber photometry recordings were performed throughout the acquisition, extinction, and reacquisition phases of trace eyeblink conditioning using an multicolor multichannel fiber photometry system (RWD R821/FR-21). Excitation light at 470 nm (GCaMP6f signal) and 410 nm (isosbestic control) was sinusoidally modulated at 15 Hz and delivered through the implanted fibers. Emitted fluorescence signals were demodulated and digitized at 1 kHz. Photometry signals were synchronized with eyelid recordings and treadmill movement using TTL pulses to ensure precise temporal alignment across behavioral and neural signals. Trials associated with the onset of locomotor rotation bouts (within 2 s) were excluded from averaged analyses based on TTL alignment.

Photometry data were analyzed offline using custom MATLAB and Python scripts. For each TTL-defined trial, fluorescence signals were extracted within an analysis window from −1.0 to 2.0 s relative to stimulus onset, with the pre-stimulus baseline window defined as −1.0 to 0.0 s. GCaMP6f and isosbestic signals were optionally smoothed using Gaussian, Savitzky–Golay, or median filtering. Baseline correction was performed using either an isosbestic fitting approach or adaptive reweighted penalized least squares (arPLS). In the isosbestic method, the 410-nm signal was fit with a biexponential function and scaled to the 470-nm signal using robust linear regression (Huber regression) to estimate motion– and bleaching-related baseline fluorescence. If fitting failed, arPLS was used as a fallback. The fitted baseline from the full recording session was interpolated to each trial and used to compute 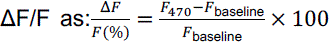. The ΔF/F trace was baseline-subtracted using the mean value of the pre-stimulus window, and z-scored signals were calculated by normalizing the mean to the baseline standard deviation. For event-related analyses, fluorescence signals were aligned to CS and US onsets. For each trial, peak Ca^2+^ responses were quantified within two post-stimulus epochs: an early window (0.0-0.32 s) and a late window (0.321-2.0 s). Only light + puff trials were included for CS– and US-aligned analyses. Heatmaps of session-by-session z-scored activity were generated for visualization. To quantify learning-dependent modulation of Ca²⁺ activity, Δz-score values were computed for CS– and US-aligned responses relative to baseline activity. Trial-by-trial Ca²⁺ responses were correlated with CR % to assess the relationship between neural activity and learning. For lateralized analysis, Ca²⁺ signals recorded from ipsilateral and contralateral cerebellar hemispheres were analyzed separately. Daily averaged Δz-scores were further computed for CS– and US-aligned responses to track the evolution of neural activity across acquisition, extinction, and reacquisition phases.

### *Ex vivo* electrophysiology

Mice were deeply anesthetized with a ketamine and xylazine mixture, and transcardially perfused with ice-cold cutting artificial cerebrospinal fluid (aCSF) containing (in mM): 87 NaCl, 2.5 KCl, 0.5 CaCl_2_, 7 MgCl_2_, 1.25 NaH_2_PO_4_, 25 NaHCO_3_, 25 glucose, and 75 sucrose, bubbled with 95% O_2_/5% CO_2_. The brain was rapidly removed and cut transversely at 300 µm using a vibratome (VT1200S, Leica Microsystems). Slices were transferred to normal aCSF containing (in mM): 119 NaCl, 2.5 KCl, 2.5 CaCl_2_, 1.3 MgCl_2_, 1.3 NaH_2_PO4, 26 NaHCO_3_, and 20 glucose, at 32°C for 30 min and then allowed to recover for 1 h at room temperature before recordings.

Whole-cell and cell-attached recordings were performed in *ex vivo* cerebellar slices prepared as described above^32, 53^. Individual slices were transferred to a submerged recording chamber mounted on a fixed-stage upright microscope (Axio Examiner.D1, Zeiss) and continuously perfused with oxygenated artificial cerebrospinal fluid (ACSF) at room temperature. PCs were visually identified using infrared differential interference contrast (IR-DIC) optics. Recordings were obtained using a MultiClamp 700B amplifier (Molecular Devices), with signals filtered at 2 kHz and digitized at 10 kHz using an ITC-18 A/D-D/A interface (Instrutech). Electrodes (3-5 MΩ) were filled with a potassium-based internal solution containing (in mM): 135 K-gluconate, 10 KCl, 10 HEPES, 1 MgCl₂, 2 Mg-ATP, and 0.3 Na-GTP (pH 7.3, 290–300 mOsm), unless otherwise indicated. Input resistance was measured using hyperpolarizing voltage steps (50 ms, 20 mV). Cells were excluded from analysis if the series resistance exceeded 25 MΩ or if any whole-cell parameter, including membrane capacitance (Cm), input resistance (Ri), or series resistance (Rs), changed by ≥20% during the recording.

Intrinsic properties were assessed in current-clamp mode. Resting membrane potential was recorded immediately after achieving whole-cell configuration. To characterize PC excitability, a series of depolarizing and hyperpolarizing current steps (typically −200 to +200 pA) were injected, and voltage responses were recorded. The number of action potentials evoked at each current step was used to generate input-output (I-O) relationships. Rheobase was defined as the minimal current required to evoke an action potential, and spike threshold was determined as the membrane potential at which the rapid upstroke of the action potential was initiated. Action potential waveform parameters were analyzed from the first spike elicited near threshold. Spike amplitude was measured from threshold to peak, and half-width was calculated at half-maximal amplitude. Fast and slow afterhyperpolarizations (fAHP and sAHP) were quantified relative to spike threshold. Additional subthreshold properties were measured, including sag amplitude during hyperpolarizing current injections, delay to first spike, and accommodation ratio, defined as the ratio of late to early inter-spike intervals during sustained firing.

To assess intrinsic excitability and plasticity, baseline recordings were performed in current-clamp mode using brief depolarizing current injections (1 s duration, 100-200 pA), adjusted for each neuron to evoke 4-10 spikes during baseline test periods. Intrinsic plasticity was induced by repetitive depolarization consisting of 150-300 pA current pulses delivered at 5 Hz for 3 s. Neuronal responses were monitored before and after induction, and changes in spike output were used to quantify intrinsic plasticity.

Spontaneous firing activity of PCs was recorded in the cell-attached configuration using aCSF-filled pipettes to form a loose seal on the cell membrane. Action potentials were detected as capacitive current transients, and firing frequency was calculated from inter-spike intervals (ISIs). Spike train variability was quantified using the coefficient of variation (CV2), defined as 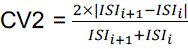, where *ISI*_i_ represents the *i*th inter-spike interval. The average CV2 across all consecutive spikes was computed for each neuron.

To evaluate synaptic input onto PCs, PFs and CFs were stimulated using aCSF-filled glass electrodes connected to an isolated stimulator (ISO-Flex, AMPI). Stimuli were delivered at 20-s intervals. Evoked excitatory postsynaptic currents (EPSCs) were recorded in voltage-clamp mode at a holding potential of –60 mV. For synaptic recordings, electrodes were filled with a cesium-based internal solution containing (in mM): 120 Cs-gluconate, 17.5 CsCl, 10 Na-HEPES, 4 Mg-ATP, 0.4 Na-GTP, 10 Na₂-creatine phosphate, and 0.2 Na-EGTA, supplemented with the sodium channel blocker QX-314 (5 mM; Tocris) (pH 7.3, 290–300 mOsm, 3-4 MΩ).

PF-PC synaptic strength was assessed by constructing input-output curves using stimulus intensities ranging from 5 to 60 µA (5, 10, 20, 30, 40, 50, and 60 µA). EPSC amplitudes were measured as peak inward currents. Paired-pulse ratios (PPRs) were measured to assess presynaptic short-term plasticity by delivering pairs of stimuli at inter-pulse intervals of 20-200 ms (in 20-ms increments), followed by 300, 400, and 500 ms intervals. PPR was calculated as the ratio of the second EPSC amplitude to the first (EPSC₂/EPSC₁). CF-PC synapses were identified by their characteristic large, all-or-none EPSCs. The threshold stimulus intensity required to evoke a CF response was first determined, and stimulus intensity was then incrementally increased to determine the number of discrete EPSC steps, reflecting CF innervation patterns onto individual PCs. For CF paired-pulse measurements, stimuli were delivered with an inter-pulse interval of 40 ms, and PPR was calculated as described above.

Spontaneous EPSCs (sEPSCs) were recorded in voltage-clamp mode at a holding potential of –60 mV in the presence of picrotoxin to block inhibitory synaptic transmission. Events were detected and analyzed using MiniAnalysis software (Synaptosoft) with a detection threshold of 8 pA. Event amplitudes and inter-event intervals were measured, and cumulative probability distributions were generated. The average sEPSC amplitude and inter-event interval were calculated for each neuron and averaged across animals.

Long-term synaptic plasticity at PF-PC synapses was induced using established stimulation protocols^61^. PF-LTD was induced by paired stimulation of PFs and CFs at 1 Hz for 5 min in current-clamp mode. Test responses were evoked at 0.05 Hz before and after induction, and EPSC amplitudes were normalized to baseline values to quantify changes in synaptic strength over time.

### Immunostaining and confocal imaging

Mice were deeply anesthetized and transcardially perfused with phosphate-buffered saline (PBS), followed by 4% paraformaldehyde in PBS. Brains were removed, post-fixed overnight at 4 °C, and sectioned in the sagittal plane (60-µm thickness) using a vibratome. Sections were stored in PBS until further processing. Free-floating sections were permeabilized in 0.25% Triton X-100 in PBS for 2 h at room temperature and blocked in 10% normal goat serum for 1 h. Sections were then incubated overnight at 4 °C with primary antibodies diluted in blocking solution.

To verify the specificity of *Mecp2* deletion in PCs, sections were co-immunostained with cell-type markers and anti-MeCP2 antibodies. Primary antibodies included mouse anti-calbindin D28K (CB; 1:500, Santa Cruz Biotechnology), rabbit anti-S100β (1:800, Sigma-Aldrich), guinea pig anti-parvalbumin (PV; 1:800, Synaptic Systems), and rabbit or mouse anti-MeCP2 (1:500, MilliporeSigma). For analysis of receptor composition and synaptic organization, sections were incubated with antibodies against AMPA receptor subunits and presynaptic markers, including rabbit anti-GluA1 (1:500, MilliporeSigma), rabbit or mouse anti-GluA2 (1:500, Thermo Fisher Scientific), rabbit anti-GluA4 (1:500, MilliporeSigma), guinea pig anti-vGluT1 (1:500, MilliporeSigma), and rabbit anti-vGluT2 (1:500, MilliporeSigma).

For morphological analysis, PC were filled with biocytin (8 mM, MilliporeSigma) during whole-cell recordings in *ex vivo* slices. Following fixation and standard processing, slices were incubated with streptavidin-conjugated Alexa Fluor 488 (1:200, Thermo Fisher Scientific) to visualize dendritic arbors. In some experiments, slices were co-immunostained with vGluT1 and vGluT2 to examine synaptic contacts along dendrites.

After primary incubation, slices were washed three times in PBS (10 min each) and incubated for 2 h at room temperature with fluorophore-conjugated secondary antibodies (1:1000; Jackson ImmunoResearch), including Alexa Fluor 488, 594, and 647 goat anti-mouse, anti-rabbit, or anti-guinea pig antibodies. Slices were washed and mounted with Vectashield mounting medium (Vector Laboratories).

Images were acquired using a Zeiss LSM 800 Airyscan confocal microscope with 10× (0.3 NA), 20× (0.8 NA), or 63× (1.4 NA oil immersion) objectives. Imaging parameters (laser power, detector gain, and pinhole size) were kept constant across groups. High-resolution z-stacks were collected for dendritic and spine analysis, while lower magnification images were used for layer-specific quantification. Image analysis was performed in ImageJ (NIH) with the experimenter blinded to genotype. Dendritic complexity was assessed using Sholl analysis, quantifying the number of dendritic intersections as a function of radial distance from the soma. Dendritic coverage area was calculated from z-projected images as the area occupied by biocytin-labeled dendrites within defined regions of interest. Spine density and volume were quantified from high-resolution images of distal dendritic segments using Imaris (Bitplane). Spine density was calculated as the number of protrusions per unit dendritic length, and spine volume was estimated based on fluorescence intensity and morphological segmentation. Presynaptic puncta (vGluT1 and vGluT2) were quantified using unbiased particle analysis in ImageJ with predefined size thresholds. Fluorescence intensity of AMPAR subunits (GluA1, GluA2, and GluA4) and presynaptic markers was measured within defined cerebellar layers and normalized across sections. CF innervation was assessed by quantifying the relative extent of vGluT2-positive puncta along PC dendrites, calculated as the ratio of the distance from the soma to the most distal vGluT2 puncta to the total dendritic length within the MCL.

### Statistics

All behavioral experiments were performed with the experimenter blinded to genotype and treatment group. Computer-assisted behavioral measurements and custom MATLAB scripts were used to further reduce bias in data acquisition and analysis. Data were analyzed using GraphPad Prism and MATLAB. Comparisons between two groups were performed using two-tailed unpaired or paired Student’s *t*-tests, Mann–Whitney two-sided tests, or Wilcoxon matched-pairs signed-rank tests, as appropriate. Repeated-measures two-way ANOVA was used for longitudinal and stimulus-response datasets, including skilled reaching, rotarod, eyeblink conditioning, Ca^2+^ imaging, Sholl analysis, intrinsic membrane responses, current-evoked spike output, input-output curves, paired-pulse analyses and synaptic plasticity measurements, with Bonferroni’s multiple-comparisons test where indicated. Fisher’s exact test was used for categorical comparisons of CF innervation pattern, Kolmogorov–Smirnov tests were used for cumulative probability analyses of mEPSCs, and simple linear regression was used for correlation analyses. Sample sizes, including the numbers of animals, cells, sections and recordings, are provided in the main text and corresponding figure legends. Data are presented as mean ± SEM. Statistical significance was defined as \**P* < 0.05, \*\**P* < 0.01 and \*\*\**P* < 0.001.

## Data Availability

Source data are provided with this paper. All data are available upon request to the corresponding author.

## Code Availability

All relevant code is available upon request to the corresponding authors.

## Acknowledgements

We are grateful to Dr. Xin Xu, Dr. Destynie Medeiros, Dr. César Acevedo-Triana, Dr. Suraj Cherian, and Dr. Akash Saxena for helpful discussions. We also thank Yijian Zhang for assistance with mouse colony maintenance. This work was supported by the United States National Institutes of Health (R21NS097913, R21NS108508, R21NS120315, and R01NS121542).

## Author Contributions

J.S., P.S., and W.L. designed the research. J.S., P.S., J.L.G., M.J., L.P., K.D., T.L., A.H.,

M.H., C.H., C.L., and W.L. conducted the research and analyzed the results. J.S., L.P.-M., and W.L. wrote the manuscript.

## Competing Interests

The authors declare no competing interests.

## Notes

### Competing Interest Statement

The authors have declared no competing interest.

