## Supplementary Figures for "MeCP2 is Necessary in Cerebellar Purkinje Cells for Precise Network Dynamics During Associative Motor Learning"

### Additional Information

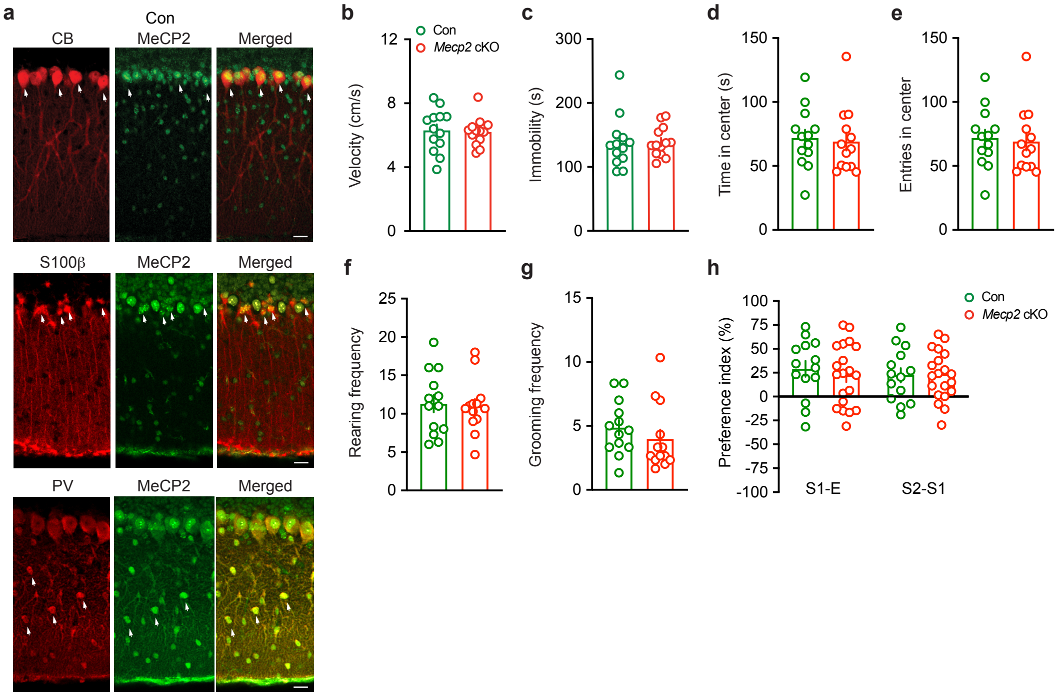


**Extended Data Fig. 1 | Validation and behavioral characterization of *Mecp2* cKO mice. a,** Immunofluorescence of cerebellar sections showing the PC marker calbindin (CB), the astrocyte marker S100β, and the interneuron marker parvalbumin (PV) with MeCP2 labeling, confirming MeCP2 expression in these cell types in control mice. Arrows indicate colocalization. Scale bar, 30 µm. **b-e,** Average travel velocity (**b**; *P* = 0.1899), immobility (**c**; *P* = 0.9404), time spent in the center (**d**; *P* = 0.7704), and number of center entries (**e**; *P* = 0.1998) during open field testing (n = 13, Con; n = 13, *Mecp2* cKO). Data are presented as mean ± SEM. Unpaired two-sided Student's *t*-test (Con vs. *Mecp2* cKO). **f,g,** Rearing (**f**) and grooming (**g**) frequencies during 10-min open field testing. Data are presented as mean ± SEM. Unpaired two-sided Student's *t*-test (n = 13, Con; n = 13, *Mecp2* cKO; *P* = 0.8130, rearing; *P* = 0.3588, grooming). **h,** Preference index (relative interaction time with the empty cup (E) and social stimuli (S1 or S2)) (n = 14, Con; n = 20,

*Mecp2* cKO). Unpaired two-sided Student's *t*-test (*P* = 0.5481, S1–E; *P* = 0.8581, S2–S1; Con vs. *Mecp2* cKO). All data are presented as mean ± SEM. Individual points represent individual animals.

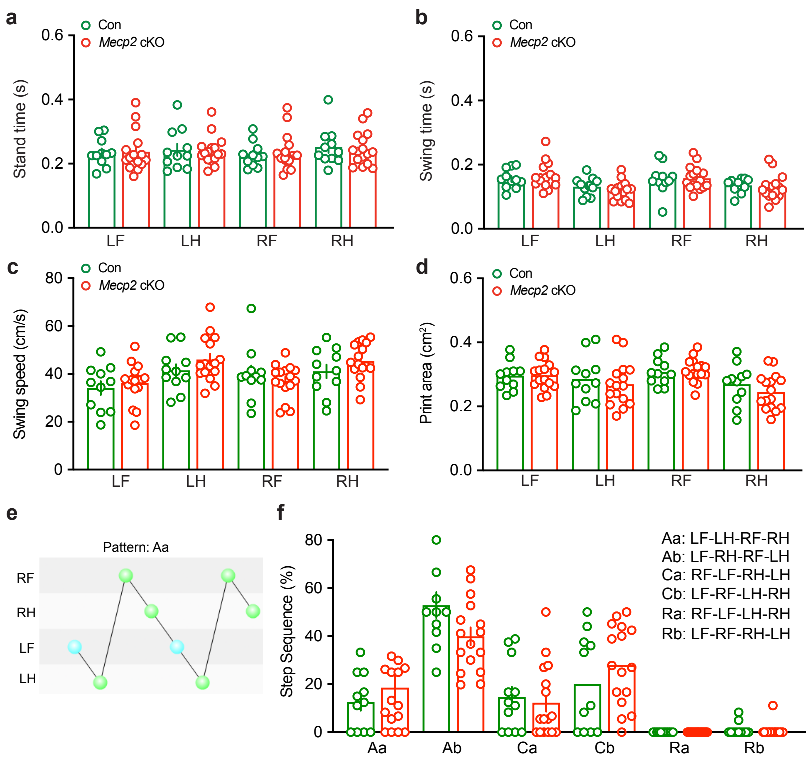


**Extended Data Fig. 2 | Additional gait parameters and step sequence analysis of *Mecp2* cKO mice. a,b,** Stand time (**a**) and swing time (**b**) for each limb (LF, LH, RF, RH) in control and *Mecp2* cKO mice. Unpaired two-sided Student's *t*-test (n = 11, Con; n = 16, *Mecp2* cKO; *P* = 0.9074, LF; *P* = 0.8785, LH; *P* = 0.6653, RF; *P* = 0.8128, RH for stand time; *P* = 0.5572, LF; *P* = 0.2717, LH; *P* = 0.8792, RF; *P* = 0.5890, RH for swing time). **c,** Swing speed for each limb (LF, LH, RF, RH). Unpaired two-sided Student's *t*-test (n = 11, Con; n = 16, *Mecp2* cKO; *P* = 0.5583, LF; *P* = 0.2111, LH; *P* = 0.3873, RF; *P* = 0.1898, RH). **d,** Print area for each limb (LF, LH, RF, RH). Unpaired two-sided Student's *t*-test (n = 11, Con; n = 16, *Mecp2* cKO; *P* = 0.9357, LF; *P* = 0.5399, LH; *P* = 0.8266, RF; *P* = 0.3191, RH). **e,** Schematic representation of step sequence patterns, with example pattern (Aa) illustrating the order of paw placement. **f,** Distribution of step sequence patterns (Aa, Ab, Ca, Cb, Ra, Rb), expressed as percentage of total steps. Unpaired two-sided Student's *t*-test or Mann–Whitney two-sided test (n = 11, Con; n = 16, *Mecp2* cKO; *P* = 0.4163, Aa; *P* = 0.0502, Ab; *P* = 0.7043, Ca; *P* = 0.2777, Cb; *P* = 0.9999, Ra; *P* = 0.5558, Rb). Step sequence definitions: Aa (LF–LH–RF–RH), Ab (LF–RH–RF–LH), Ca (RF–LF–RH–LH), Cb (LF–RF–LH–RH), Ra (RF–LF–LH–RH), and Rb (LF–RF–RH–LH). All data are presented as mean ± SEM. Individual points represent individual animals.

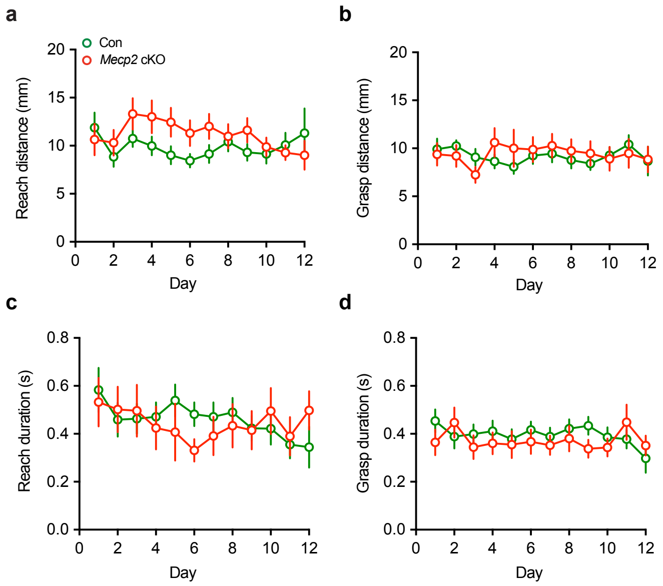


**Extended Data Fig. 3 | Additional skilled reaching performance analysis in *Mecp2* cKO mice**. **a,** Reach distance across training days. Two-way repeated-measures ANOVA showed no day × genotype interaction $F(11,383) = 1.361$, $P = 0.1890$, no main effect of day $F(11,383) = 1.500$, $P = 0.1287$), and no main effect of genotype $F(1,51) = 1.619$, $P = 0.2090$ (n = 27, Con; n = 26, *Mecp2* cKO). **b,** Grasp distance across training days. Two-way repeated-measures ANOVA showed no day × genotype interaction $F(11,383) = 0.8722$, $P = 0.5679$, no main effect of day $F(11,383) = 0.4771$, $P = 0.9172$), and no main effect of genotype $F(1,51) = 0.03014$, $P = 0.8629$ (n = 27, Con; n = 26, *Mecp2* cKO). **c,** Reach time across training days. Two-way repeated-measures ANOVA showed no day × genotype interaction $F(13,397) = 0.5190$, $P = 0.9130$, no main effect of day $F(13,397) = 0.6822$, $P = 0.7811$), and no main effect of genotype $F(1,51) = 0.09032$, $P = 0.7650$ (n = 25, Con; n = 28, *Mecp2* cKO). **d,** Grasp time across training days. Two-way repeated-measures ANOVA showed no day × genotype interaction $F(13,397) = 1.414$, $P = 0.1497$, no main effect of day $F(13,397) = 0.9018$, $P = 0.5513$), and no main effect of genotype

$F(1,51) = 0.2678$, $P = 0.6071$ (n = 25, Con; n = 28, *Mecp2* cKO). All data are presented as mean ± SEM.

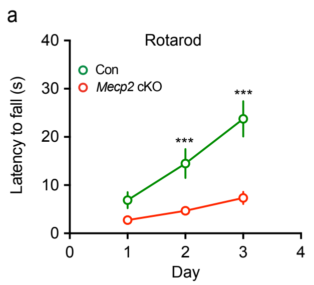


**Extended Data Fig. 4 | Impaired rotarod performance in *Mecp2* cKO mice. a,** Latency to fall during rotarod training in control and *Mecp2* cKO mice. Mice underwent 3 consecutive days of rotarod training, and latency to fall was averaged across trials for each day. Two-way repeated-measures ANOVA showed a significant day × genotype interaction $F(2, 52) = 8.686$, $P < 0.001$, a significant main effect of day $F(2, 52) = 26.71$, $P < 0.001$, and a significant main effect of genotype $F(1, 26) = 27.67$, $P < 0.001$ (n = 11, Con; n = 17, *Mecp2* cKO). All data are presented as mean ± SEM.

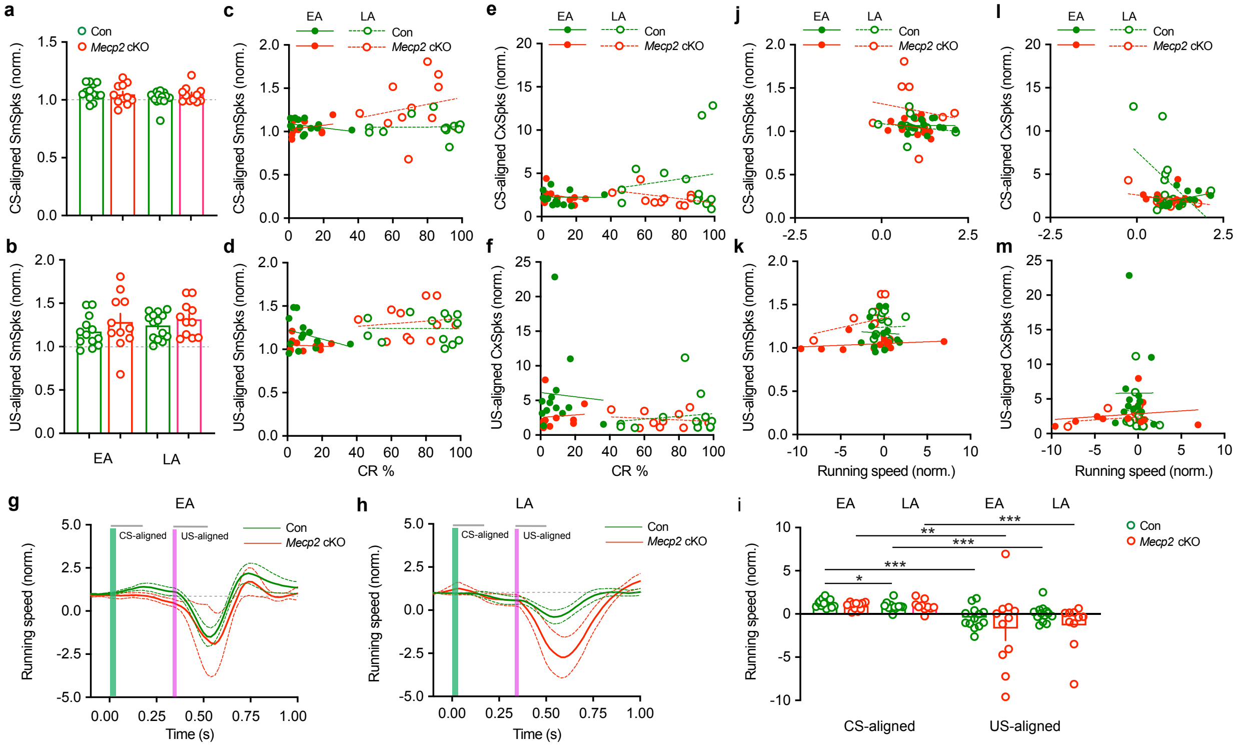


**Extended Data Fig. 5 | Relationship between PC firing, CRs, and locomotion during trace eyeblink conditioning in *Mecp2* cKO mice. a,b,** Quantification of CS-aligned normalized SmSpk responses (**a**) and US-aligned normalized SmSpk responses (**b**) in control and *Mecp2* cKO mice during EA and LA. Wilcoxon matched-pairs signed-rank test showed no significant difference between EA and LA in CS-aligned SmSpk responses (**a**; n = 13, Con; n = 11, *Mecp2* cKO; *P* = 0.1272, Con EA vs. Con LA; *P* > 0.9999, *Mecp2* cKO EA vs. *Mecp2* cKO LA) and US-aligned normalized SmSpk responses (**b**; n = 13, Con; n = 11, *Mecp2* cKO; *P* = 0.3757, Con EA vs. Con LA; *P* = 0.7646, *Mecp2* cKO EA vs. *Mecp2* cKO LA). **c-f,** Correlation of CS-aligned normalized SmSpk responses (**c**), US-aligned normalized SmSpk responses (**d**), CS-aligned normalized CxSpk responses (**e**), and US-aligned normalized CxSpk responses (**f**) with CR percentage during EA and LA in control and *Mecp2* cKO mice. Spike counts were calculated in 20-ms bins, aligned to

CS and US onset, and normalized to baseline activity. Simple linear regression showed no significant correlation between CR percentage and CS-aligned SmSpk responses (**c**; n = 13, Con; n = 11, *Mecp2* cKO; WT EA: $R^2 = 0.1089$, $F(1,11) = 1.344$, $P = 0.2709$; *Mecp2* cKO EA: $R^2 = 0.06232$, $F(1,9) = 0.5982$, $P = 0.4591$; WT LA: $R^2 = 4.411 \times 10^{-5}$, $F(1,11) = 0.0004852$, $P = 0.9828$; *Mecp2* cKO LA: $R^2 = 0.04018$, $F(1,9) = 0.3767$, $P = 0.5546$), US-aligned SmSpk responses (**d**; n = 13, Con; n = 11, *Mecp2* cKO; WT EA: R2 = 0.08478, $F(1,11) = 1.019$, $P = 0.3344$; *Mecp2* cKO EA: $R^2 = 0.01004$, $F(1,9) = 0.09129$, $P = 0.7694$; WT LA: $R^2 = 0.0001132$, $F(1,11) = 0.001245$, $P = 0.9725$; *Mecp2* cKO LA: $R^2 = 0.01518$, $F(1,9) = 0.1387$, $P = 0.7182$), CS-aligned CxSpk responses (**e**; n = 13, Con; n = 11, *Mecp2* cKO; WT EA: $R^2 = 0.0003265$, $F(1,11) = 0.003593$, $P = 0.9533$; *Mecp2* cKO EA: $R^2 = 0.1054$, $F(1,9) = 1.061$, $P = 0.3300$; WT LA: $R^2 = 0.02882$, $F(1,11) = 0.3262$, $P = 0.6223$; *Mecp2* cKO LA: $R^2 = 0.2176$, $F(1,9) = 2.503$, $P = 0.1481$), and US-aligned CxSpk responses (**f**; n = 13, Con; n = 10, *Mecp2* cKO; WT EA: $R^2 = 0.002684$, $F(1,11) = 0.02960$, $P = 0.8665$; *Mecp2* cKO EA: $R^2 = 0.005993$, $F(1,8) = 0.04823$, $P = 0.8317$; WT LA: $R^2 = 0.01941$, $F(1,11) = 0.2177$, $P = 0.6499$; *Mecp2* cKO LA: $R^2 = 0.02528$, $F(1,8) = 0.2075$, $P = 0.6608$). **g,h,** Averaged running speed profiles aligned to CS and US onset during EA (**g**) and LA (**h**) in control and *Mecp2* cKO mice. Treadmill movement was synchronized with electrophysiological recordings using TTL pulses. **i,** Quantification of normalized running speed during CS-aligned and US-aligned epochs in control and *Mecp2* cKO mice during EA and LA. Mann-Whitney two-sided test showed significant differences between conditions (n = 13, Con EA; n = 11, *Mecp2* cKO EA; n = 13, Con LA; n = 10, *Mecp2* cKO LA; $P = 0.0441$, CS-aligned WT EA vs. WT LA; $P < 0.001$, CS-aligned WT EA vs. US-aligned WT EA; $P < 0.001$, CS-aligned WT LA vs. US-aligned WT LA; $P <$

0.001, CS-aligned *Mecp2* cKO EA vs. US-aligned *Mecp2* cKO EA; $P < 0.001$, CS-aligned *Mecp2* cKO LA vs. US-aligned *Mecp2* cKO LA). **j-m,** Correlation of normalized running speed with CS-aligned normalized SmSpk responses (**j**), CS-aligned normalized SmSpk responses (**k**), US-aligned normalized SmSpk responses (**l**), and US-aligned normalized CxSpk responses (**m**) in control and *Mecp2* cKO mice. Simple linear regression showed no significant correlation between running speed and CS-aligned SmSpk responses (**j**; n = 13, Con EA; n = 11, *Mecp2* cKO EA; n = 13, Con LA; n = 10, *Mecp2* cKO LA; WT EA: $R^2 = 4.661 \times 10^{-5}$, $F(1,11) = 0.0005128$, $P = 0.9823$; *Mecp2* cKO EA: $R^2 = 0.07712$, $F(1,9) = 0.7521$, $P = 0.4084$; WT LA: $R^2 = 0.03312$, $F(1,11) = 0.3768$, $P = 0.5518$; *Mecp2* cKO LA: $R^2 = 0.02644$, $F(1,8) = 0.2173$, $P = 0.6536$), US-aligned SmSpk responses (**k**; n = 13, Con EA; n = 11, *Mecp2* cKO EA; n = 13, Con LA; n = 10, *Mecp2* cKO LA; WT EA: $R^2 = 0.001878$, $F(1,11) = 0.02069$, $P = 0.8882$; *Mecp2* cKO EA: $R^2 = 0.06946$, $F(1,9) = 0.6718$, $P = 0.4336$; WT LA: $R^2 = 0.0006408$, $F(1,11) = 0.007054$, $P = 0.9346$; *Mecp2* cKO LA: $R^2 = 0.08061$, $F(1,8) = 0.7014$, $P = 0.4266$), CS-aligned CxSpk responses (**l**; n = 13, Con EA; n = 11, *Mecp2* cKO EA; n = 13, Con LA; n = 10, *Mecp2* cKO LA; WT EA: $R^2 = 0.1962$, $F(1,11) = 2.685$, $P = 0.1296$; *Mecp2* cKO EA: $R^2 = 0.003153$, $F(1,9) = 0.02847$, $P = 0.8697$; WT LA: $R^2 = 0.2343$, $F(1,11) = 3.366$, $P = 0.0937$; *Mecp2* cKO LA: $R^2 = 0.1454$, $F(1,8) = 1.361$, $P = 0.2770$), and US-aligned CxSpk responses (**m**; n = 13, Con EA; n = 10, *Mecp2* cKO EA; n = 13, Con LA; n = 10, *Mecp2* cKO LA; WT EA: $R^2 = 6.559 \times 10^{-5}$, $F(1,11) = 0.0006559$, $P = 0.9937$; *Mecp2* cKO EA: $R^2 = 0.03624$, $F(1,8) = 0.3008$, $P = 0.5983$; WT LA: $R^2 = 0.03279$, $F(1,11) = 0.3729$, $P = 0.5538$; *Mecp2* cKO LA: $R^2 = 0.03574$, $F(1,8) = 0.2965$, $P = 0.6009$). All data are presented as mean ± SEM. Individual points represent individual animals.

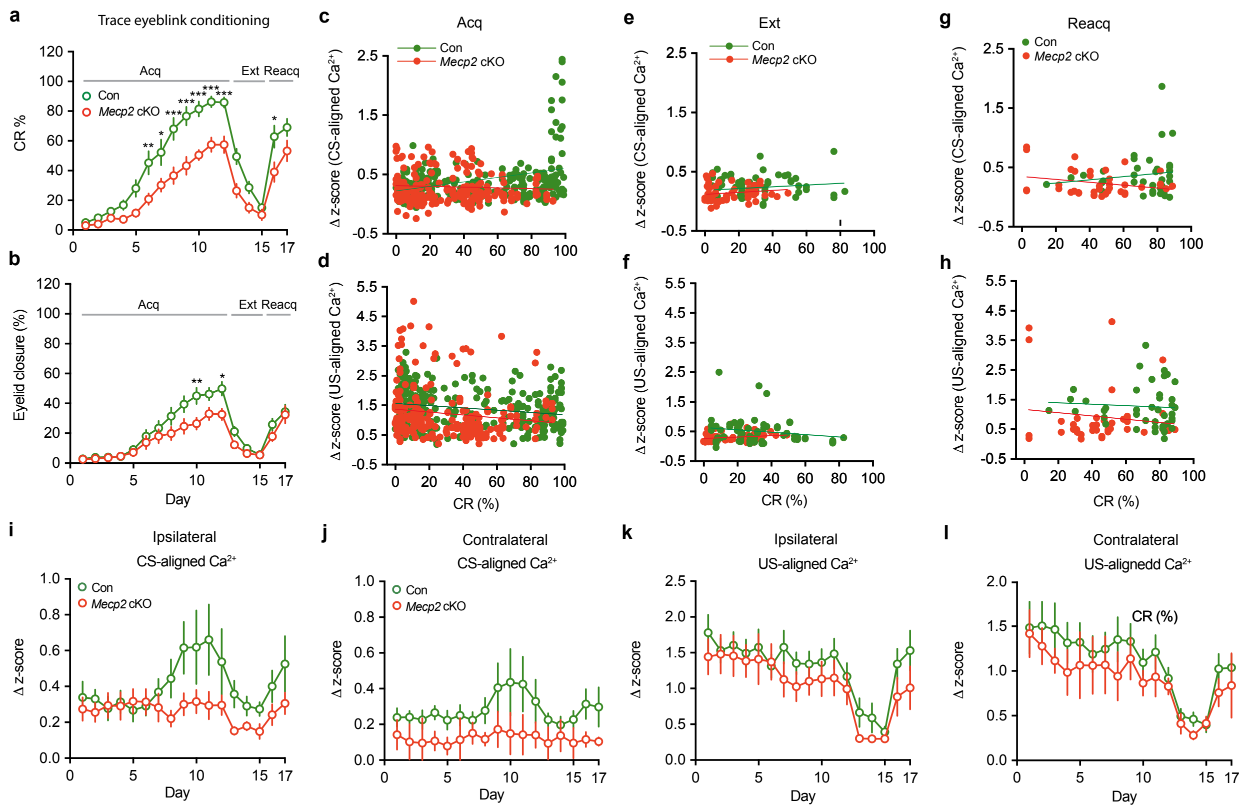


**Extended Data Fig. 6 | Relationship between PC $Ca^{2+}$ signals and behavioral performance during trace eyeblink conditioning in *Mecp2* cKO mice. a,b,** Quantification of CR percentage (**a**) and eyelid closure percentage (**b**) across acquisition (Acq), extinction (Ext), and reacquisition (Reacq) during trace eyeblink conditioning in control and *Mecp2* cKO mice. **c-h,** Relationship between CS-aligned $Ca^{2+}$ responses (**c,e,g**) or US-aligned $Ca^{2+}$ responses (**d,f,h**) and CR percentage during acquisition (**c,d**), extinction (**e,f**), and reacquisition (**g,h**) in control and *Mecp2* cKO mice. Each point represents a session-averaged $Ca^{2+}$ response plotted against behavioral performance. $Ca^{2+}$ signals were extracted within an analysis window from -1.0 to 2.0 s relative to stimulus onset, baseline-subtracted, and expressed as z-scored dF/F. Simple linear regression showed a significant correlation between CR percentage and CS-aligned $Ca^{2+}$

responses during acquisition (**c**; n = 252, Con; n = 228, *Mecp2 cKO*; WT: $R^2$ = 0.1264, $F(1,250)$ = 36.17, $P < 0.001$; *Mecp2* cKO: $R^2$ = 0.002382, $F(1,226)$ = 0.5396, $P$ = 0.4634), and between CR percentage and US-aligned $Ca^{2+}$ responses during acquisition (**d**; n = 252, Con; n = 228, *Mecp2* cKO; WT: $R^2$ = 0.03878, $F(1,250)$ = 10.09, $P$ = 0.0017; *Mecp2* cKO: $R^2$ = 0.01877, $F(1,226)$ = 4.323, $P$ = 0.0387). No significant correlations were detected between CR percentage and CS-aligned $Ca^{2+}$ responses during extinction (**e**; n = 60, Con; n = 57, *Mecp2* cKO; WT: $R^2$ = 0.0002652, $F(1,58)$ = 0.01539, $P$ = 0.9017; *Mecp2* cKO: $R^2$ = 0.003466, $F(1,55)$ = 0.1913, $P$ = 0.6635) or reacquisition (**g**; n = 38, Con; n = 38, *Mecp2* cKO; WT: $R^2$ = 0.01723, $F(1,36)$ = 0.6313, $P$ = 0.4321; *Mecp2* cKO: $R^2$ = 0.01477, $F(1,36)$ = 2.504, $P$ = 0.1223), or between CR percentage and US-aligned $Ca^{2+}$ responses during extinction (**f**; n = 63, Con; n = 57, *Mecp2* cKO; WT: $R^2$ = 0.03049, $F(1,61)$ = 1.918, $P$ = 0.1711; *Mecp2* cKO: $R^2$ = 0.02674, $F(1,57)$ = 1.511, $P$ = 0.2242) or reacquisition (**h**; n = 42, Con; n = 38, *Mecp2* cKO; WT: $R^2$ = 0.003162, $F(1,40)$ = 0.1269, $P$ = 0.7236; *Mecp2* cKO: $R^2$ = 0.02674, $F(1,36)$ = 0.5397, $P$ = 0.4673). **i-l**, Quantification of ipsilateral and contralateral CS-aligned $Ca^{2+}$ responses (**i,j**) and US-aligned $Ca^{2+}$ responses (**k,l**) across acquisition, extinction, and reacquisition in control and *Mecp2* cKO mice. For ipsilateral CS-aligned $Ca^{2+}$ responses (**i**), two-way repeated-measures ANOVA showed a significant day × genotype interaction $F(16,384)$ = 1.914, $P$ = 0.0180), a significant main effect of day $F(16,384)$ = 3.248, $P < 0.001$), and a trend toward significant main effect of genotype $F(1,24)$ = 3.017, $P$ = 0.0952) (n = 11, Con; n = 15, *Mecp2* cKO). For contralateral CS-aligned $Ca^{2+}$ responses (**j**), two-way repeated-measures ANOVA showed no significant day × genotype interaction $F(16,192)$ = 0.2647, $P$ = 0.9982, no significant main effect of day $F(16,192)$ = 0.5666, $P$ = 0.9059, and a trend toward significant main effect of genotype $F(1,12) = 2.674$, $P = 0.1279$ (n = 10, Con; n = 4, *Mecp2* cKO). For ipsilateral US-aligned $Ca^{2+}$ responses (**k**), two-way repeated-measures ANOVA showed no significant day × genotype interaction $F(16,384) = 0.6808$, $P = 0.8134$), a significant main effect of day $F(16,384) = 16.58$, $P < 0.001$, and no significant main effect of genotype $F(1,24) = 0.8870$, $P = 0.3557$) (n = 11, Con; n = 15, *Mecp2* cKO). For contralateral US-aligned $Ca^{2+}$ responses (**l**), two-way repeated-measures ANOVA showed no significant day × genotype interaction $F(16,192) = 0.2439$, $P = 0.9989$), a significant main effect of day $F(16,192) = 8.551$, $P < 0.001$), and no significant main effect of genotype $F(1,12) = 0.5476$, $P = 0.4735$) (n = 10, Con; n = 4, *Mecp2* cKO). All data are presented as mean ± SEM. Individual points represent individual animals.

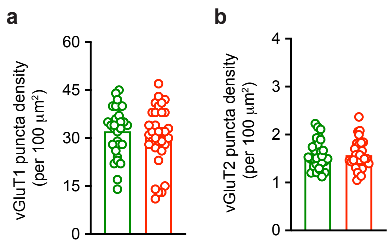


**Extended Data Fig. 7 | vGluT1 and vGluT2 puncta density in control and *Mecp2* cKO mice. a,** Quantification of vGluT1 puncta density. Unpaired two-sided Student's *t*-test (n = 31, Con; n = 30, *Mecp2* cKO; *P* = 0.8738). **b,** Quantification of vGluT2 puncta density. Unpaired two-sided Student's *t*-test (n = 30, Con; n = 33, *Mecp2* cKO; *P* = 0.4918). All data are presented as mean ± SEM. Individual points represent individual sections.

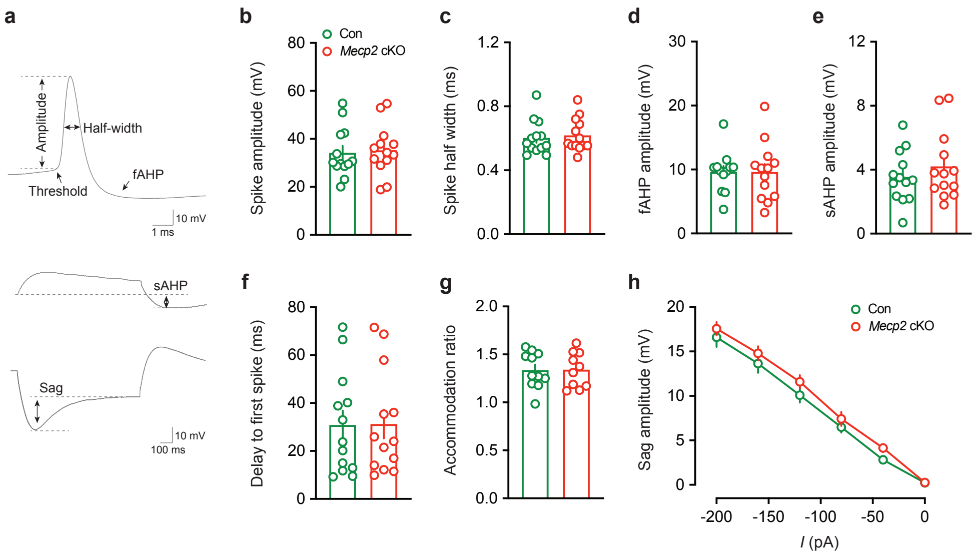


**Extended Data Fig. 8 | Passive and active membrane properties of PCs in *Mecp2* cKO mice. a,** Schematic illustrating the measurements of action potential amplitude, spike half-width, fast afterhyperpolarization (fAHP), slow afterhyperpolarization (sAHP), and sag amplitude. **b-g,** Quantification of spike amplitude (**b**), spike half-width (**c**), fAHP amplitude (**d**), sAHP amplitude (**e**), delay to first spike (**f**), and accommodation ratio (**g**). Unpaired two-sided Student's *t*-test (n = 13, Con; n = 13, *Mecp2* cKO; $P = 0.7872$, spike amplitude; $P = 0.6968$, spike half-width; $P = 0.9697$, fAHP amplitude; $P = 0.3865$, sAHP amplitude; $P = 0.9704$, delay to first spike; $P = 0.9988$, accommodation ratio). **h,** Quantification of membrane voltage sag amplitude across hyperpolarizing current injections. Two-way repeated-measures ANOVA with Bonferroni's multiple-comparisons test showed a significant current × genotype interaction $F(7, 189) = 2.166$, $P = 0.0390$, a significant main effect of input current $F(7, 189) = 540.7$, $P < 0.001$, and no significant main effect of genotype $F(1, 27) = 0.05661$, $P = 0.8137$ (n = 16, Con; n = 14, *Mecp2* cKO). All data are presented as mean ± SEM. Individual points represent individual recordings.

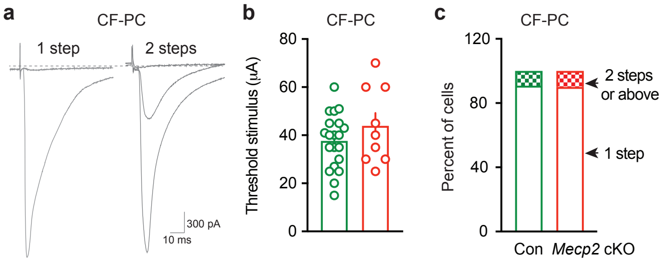


**Extended Data Fig. 9 | Preserved climbing fiber–Purkinje cell synaptic innervation in *Mecp2* cKO mice. a,** Representative CF-PC EPSCs showing one-step and two-step CF responses. **b,** Quantification of the threshold stimulus intensity required to evoke CF-PC EPSCs. Unpaired two-sided Student's *t*-test (n = 20, Con; n = 9, *Mecp2* cKO; *P* = 0.2221). **c,** Quantification of the proportion of PCs exhibiting 1-step versus 2-step-or-above CF innervation. Fisher's exact test showed no significant difference between groups (Con: 18 cells with 1-step and 2 cells with 2-step-or-above CF innervation; *Mecp2* cKO: 9 cells with 1-step and 1 cell with 2-step-or-above CF innervation; *P* > 0.9999. All data are presented as mean ± SEM. Individual points represent individual recordings.
